# Sub-neutralizing concentrations of Zika virus IgM monoclonal antibodies reduce IgG-mediated antibody-dependent enhancement of infectivity

**DOI:** 10.64898/2026.01.03.697490

**Authors:** Matthias Lingemann, Taylor J. McGee, Giacomo Sidoti Migliore, Jenna M. DeLuca, Stephen S. Whitehead, Mattia Bonsignori

**Author notes:** These authors contributed equally to this work (Matthias Lingemann, Taylor J. McGee, Giacomo Sidoti Migliore).

## Abstract

Zika virus (ZIKV) is an arthropod-borne flavivirus endemic to Latin America and Africa for which IgG-mediated antibody-dependent enhancement of infectivity (ADE) is a phenomenon of concern. The interplay dynamics between flavivirus-specific antibodies of the IgG and IgM isotypes are mostly unknown. Here, we used DH1017.IgM, a previously described human-derived, pentameric, ultrapotent neutralizing ZIKV IgM monoclonal antibody (mAb) and a recombinantly produced IgM version of neutralizing EDE1 mAb C8 (C8.IgM) to assess if IgM antibodies can interfere with IgG-mediated ADE *in vitro*. Both mAbs consistently reduced IgG-mediated ADE across a panel of ZIKV-binding IgG mAbs in a dose-dependent manner. ADE mitigation was achieved at DH1017.IgM sub-neutralizing concentrations. C8.IgM also mitigated IgG-mediated ADE against the whole panel, demonstrating generalizability. Finally, DH1017.IgM confirmed inhibition of ADE at sub-neutralizing concentrations in sera of ZIKV-infected individuals. In conclusion, mAbs of the IgM isotype may potently counter the adverse effects of IgG-mediated ADE.

## INTRODUCTION

Zika virus (ZIKV) and dengue virus (DENV) co-circulate in vast endemic areas. While antibody-dependent enhancement of infectivity (ADE) is a well-documented issue in DENV infections, where it can be associated with severe disease, and has been invoked as the cause of enhanced disease in seronegative recipients of a tetravalent live-attenuated chimeric DENV vaccine,^1–3^ the clinical significance of ADE in ZIKV infection remains uncertain. Nonetheless, there is ample evidence that ZIKV infectivity can be enhanced *in vitro* by antibodies of the IgG isotype: convalescent plasma from DENV-infected individuals can mediate a substantial increase in ZIKV infection *in vitro*;^4–7^ ZIKV-immune plasma from both symptomatic and asymptomatic individuals, as well as ZIKV and DENV monoclonal antibodies (mAb), can sustain ZIKV ADE both *in vitro* and in mice;^8,9^ and finally, *in vivo* studies in pregnant non-human primates and mice, as well as studies on *ex vivo* human placentas, suggest that ADE may contribute to ZIKV placental pathogenesis and favor ZIKV vertical transmission.^8,10–12^ The potential role of ADE in pregnancy is particularly problematic because ZIKV infection in pregnancy carries the greatest burden of disease due to the high frequency of severe and often lethal fetal neurological birth defects, including microcephaly, cumulatively known as Congenital Zika Syndrome (CZS).^13–19^ Therefore, ADE in ZIKV infections remains a phenomenon of concern.

The underlying mechanism of ADE has been imputed to flaviviruses exhibiting a remarkable resistance to lysis by monocytes and macrophages, and to their ability to escape and replicate in these cells, where they are chaperoned through IgG opsonization.^20^ Hence, IgG antibodies can mediate both neutralization and ADE as a function of their affinity, neutralization potency, concentration, and stoichiometry of virion binding.^21,22^

In contrast, antibodies of the IgM isotype are not known to mediate ADE. The immunobiology of ZIKV humoral responses, similarly to other flavivirus infections,^23–27^ is characterized by prolonged serum ZIKV IgM titers that can persist from months to years and contribute to ZIKV neutralization.^28–31^ Importantly, this phenomenon also applies to ZIKV infections during pregnancy, and we demonstrated that neutralizing IgM can contribute to ZIKV serum neutralization in pregnancy for up to three months after maternal development of Zika symptoms.^30^

A previous study using IgG- and IgM-depleted pooled sera from human primary ZIKV infections indicated that early ZIKV serum IgM may counteract IgG-mediated ADE and it was speculated that ADE mitigation may be due to increased steric hindrance of IgM in preventing less avid IgG antibodies to bind.^32^ However, the dynamics of IgM and IgG combinatorial activities are likely more complex and have not been further characterized. To fill this knowledge gap, interactions of native pentameric IgM and IgG mAbs need to be investigated.

We had previously characterized ZIKV ultrapotent neutralizing IgM mAb DH1017.IgM in its native pentameric conformation.^30^ DH1017.IgM was isolated at 71 days post symptom onset from a pregnant woman with Zika who gave birth to a healthy baby. DH1017.IgM binds to a ZIKV E-dimer quaternary and discontinuous epitope encompassing Domain II (DII) and the DI/DII interface. Its ultrapotent neutralization depends upon its multimeric structure. We identified two non-mutually exclusive modes of multivalent virion recognition: (1) the pentameric IgM molecule lays parallel to the virion surface, assumes an umbrella-like conformation and engages distinct ZIKV E-dimers on the same virion, or (2) the pentameric IgM in its planar conformation assumes a perpendicular orientation relative to the virion surface and cross-links multiple virions.^30^ We estimated that only three pentameric IgM can bind concurrently to the same virion in the umbrella-like conformation, occupying only one third of the cognate epitopes exposed on the virion surface.^30^ As expected, DH1017.IgM did not mediate *in vitro* ADE in primary monocytes, human monocytic THP-1 cells, and K562 cells.^30^ Due to its functional characteristics, we had proposed IgM-based biologics as potential countermeasures against ZIKV infections, especially to prevent vertical transmission.

Here, we use DH1017.IgM as a prototypic ZIKV IgM mAb to evaluate the combinatorial effect of DH1017.IgM and a panel of ZIKV IgG mAbs on multiple antibody functions, including virion binding, neutralization, and ADE. We illustrate the complex dynamics of IgG/IgM cross-competition for binding to ZIKV whole virions, which lead to incomplete reciprocal blocking; we demonstrate that the cross-competition between DH1017.IgM and ZIKV IgG mAbs does not exert negative effects on viral neutralization, and we provide direct evidence that DH1017.IgM reduced IgG mAb-mediated ADE in a dose-dependent manner and up to full abrogation, regardless of IgG epitope specificity. Remarkably, DH1017.IgM exerts ADE reduction at sub-neutralizing concentrations. That such reduction could be observed with a different recombinantly-produced IgM mAb (C8.IgM) demonstrates that this is not a function exclusive to DH1017.IgM. Thus, our study indicates that IgM ADE reduction is a function distinct from viral neutralization and likely multifactorial.

We further show that DH1017.IgM mAb can reduce *in vitro* IgG-mediated ADE in unfractionated sera of primary ZIKV infections. These findings suggest that DH1017.IgM might provide protective benefits beyond neutralization -i.e., exceptionally potent ADE reduction through passive immunization and further support the potential use of multivalent IgM-based biologics as countermeasures against ZIKV infection and vertical transmission.

## RESULTS

### ZIKV virion binding cross-competition between DH1017.IgM and ZIKV IgG mAbs is incomplete

Given the DH1017.IgM large footprint and the associated steric hindrance,^30^ we analyzed how DH1017.IgM and other ZIKV mAbs of the IgG isotype reciprocally affect binding to ZIKV/Aedes-africanus/SEN/DakAr41524/1984 whole virion (ZIKV-DAK) in a competitive environment. To do this, we produced a panel of 14 well-characterized IgG mAbs targeting multiple regions of the ZIKV E glycoprotein (**Table 1**). Each IgG mAb and recombinantly produced pentameric DH1017.IgM mAb were directly conjugated to HRP for competition ELISA. The ability of each mAb to compete its own HRP-conjugated version was preliminarily confirmed (**Figure S1**). DH1017.IgM competed autologous binding (**Figure S1A)**, as well as binding of its IgG version (DH1017.IgG), which more closely reflected the experimental assay condition (**Figure S1B**). Similarly, of the 14 ZIKV-IgG mAbs in the panel, 12 competed binding of their respective HRP-conjugated version (**Figure S1C-N**) and were used in reciprocal binding competition ELISA with DH1017.IgM. DII Z20 and DIII Z23 mAbs failed to self-compete and were excluded from cross-competition assays.

**Table 1:**
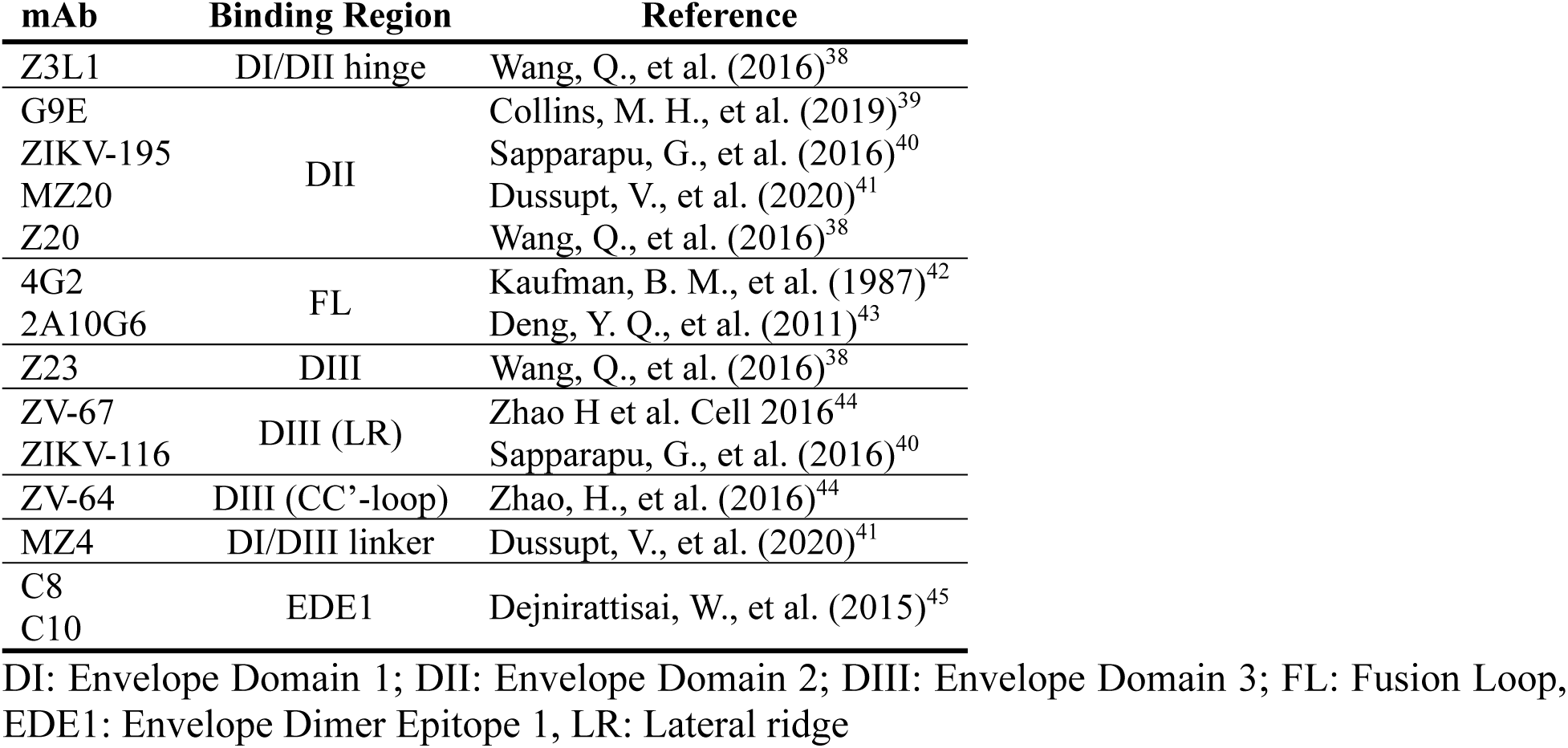
ZIKV-reactive IgG mAb panel.

| **mAb** | **Binding Region** | **Reference** |
| --- | --- | --- |
| Z3L1 | DI/DII hinge | Wang, Q., et al. (2016)<sup>38</sup> |
| G9E | DII | Collins, M. H., et al. (2019)<sup>39</sup> |
| ZIKV-195 |  | Sapparapu, G., et al. (2016)<sup>40</sup> |
| MZ20 |  | Dussupt, V., et al. (2020)<sup>41</sup> |
| Z20 |  | Wang, Q., et al. (2016)<sup>38</sup> |
| 4G2 | FL | Kaufman, B. M., et al. (1987)<sup>42</sup> |
| 2A10G6 |  | Deng, Y. Q., et al. (2011)<sup>43</sup> |
| Z23 | DIII | Wang, Q., et al. (2016)<sup>38</sup> |
| ZV-67 | DIII (LR) | Zhao H et al. Cell 2016<sup>44</sup> |
| ZIKV-116 |  | Sapparapu, G., et al. (2016)<sup>40</sup> |
| ZV-64 | DIII (CC'-loop) | Zhao, H., et al. (2016)<sup>44</sup> |
| MZ4 | DI/DIII linker | Dussupt, V., et al. (2020)<sup>41</sup> |
| C8 | EDE1 | Dejnirattisai, W., et al. (2015)<sup>45</sup> |
| C10 |  |  |

DI: Envelope Domain 1; DII: Envelope Domain 2; DIII: Envelope Domain 3; FL: Fusion Loop, EDE1: Envelope Dimer Epitope 1, LR: Lateral ridge

DIII region ZV-67, ZIKV-116 and ZV-64 and DI/DIII linker region MZ4 mAbs did not cross compete with DH1017.IgM (IC_50_ >100 nM) (**Figure 1A**), which is in line with the epitope footprint of DH1017.IgM not including the DIII region.^30^ While DH1017.IgM was outcompeted by EDE1 C10 (DH1017.IgM IC_50_ = 39 nM vs C10 IC_50_ = 1.7 nM) and DII region MZ20 mAbs (DH1017.IgM IC_50_ >100 nM vs MZ20 IC_50_ = 2.3 nM), DH1017.IgM outcompeted the remaining six IgG mAbs, representing all the remaining ZIKV E glycoprotein epitope regions included in the panel (**Figure 1A**). However, while DH1017.IgM competed most ZIKV IgG mAbs more potently than vice versa, it was less efficient than ZIKV IgGs in achieving complete blocking. At saturating concentrations, half of the IgG mAbs in the panel (6 ot of 12) competed >75% DH1017.IgM binding (**Figure 1B** and **S2**), whereas DH1017.IgM vastly failed to efficiently block IgG binding: except for three mAbs (DI/DII hinge Z3L1, DII ZIKV-195 and EDE1 C8), DH1017.IgM blocked <75% of IgG binding (**Figure 1C** and **S3**).

**Figure 1:**
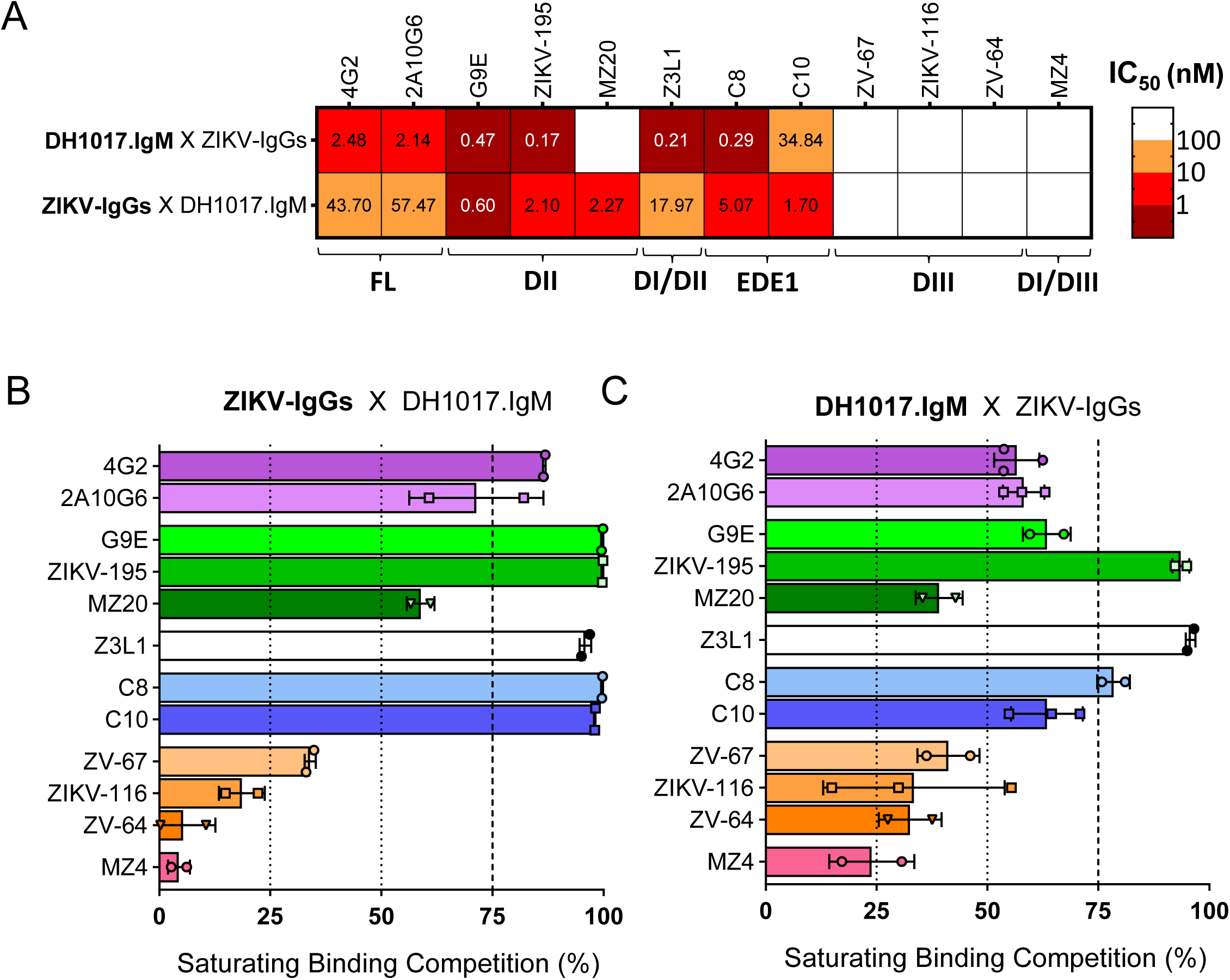
ZIKV-DAK whole virion binding cross-competition of DH1017.IgM and ZIKV-IgG mAbs. **(A)** Heatmap analysis of ZIKV-DAK whole virion binding competition between DH1017.IgM and 12 ZIKV-IgG mAbs binding (top) and vice versa (bottom). Coloration is by IC_50_. Binding regions of each ZIKV-IgG mAb are indicated as shown in Table 1. **(B,C)** Percentage ZIKV IgG mAb competition of DH1017.IgM binding (B) and vice versa (C) to ZIKV-DAK whole virion. Blocking mAbs were used at saturating concentration of 100 µg/ml. Each bars shows the average of two independent experiments and is color-coded based on each IgG binding region (purple: fusion loop; green: DII; white: DI/DII hinge; blue: EDE1; orange: DIII; pink: DI/DIII linker region. Error bars: SD of the two experimental replicates. Each experiment was performed in technical duplicates. See also Figures S1, S2 and S3.

These findings demonstrate that, at equimolar concentrations, DH1017.IgM binding to ZIKV is unlikely to be blocked by the presence of ZIKV IgG antibodies in a polyclonal environment. However, they critically indicate that, even at full DH1017.IgM occupancy, a fraction of the epitopes expressed on the virion surface remain available for engagement by IgGs directed against multiple E glycoprotein domains and regions, suggesting that DH1017.IgM may influence the binding stoichiometry of other ZIKV-specific IgGs on the virion.

### Combinations of ZIKV IgG mAbs and DH1017.IgM retain their respective neutralizing capacities

ADE can be mediated by both non-neutralizing and neutralizing IgG antibodies, and IgG binding stoichiometry is a critical factor for ADE.^21^ The relationship between the concentrations at which a neutralizing antibody mediates peak ADE and neutralization is theoretically important. If peak ADE concentrations are much lower than neutralizing concentrations, but still physiologically relevant (e.g., in the waning phases of humoral responses post-acute infection, or at steady state after infection due to long-lived plasma cells), there is a potential increased risk of *in vivo* ADE.^20,21^ Therefore, we focused our attention on the effect of DH1017.IgM co-incubation with ZIKV IgG mAbs on neutralization and ADE activity.

We preliminarily assessed ADE and neutralizing activities of 14 ZIKV IgG mAbs against reporter virus particles (RVP) expressing the C-prM-E proteins of ZIKV-DAK. Apart from DIII ZV-64, all ZIKV IgG mAbs mediated measurable ADE (**Figure S4**) and with the exception of FL 4G2 and 2A10G6, DII Z20, and DIII ZV-64, all IgG mAbs neutralized 50% of ZIKV-DAK infectivity at concentrations <100 µg/ml (**Figure S5**). DIII ZV-64 was excluded from further analyses due to concurrent lack of ADE and neutralizing activity against ZIKV-DAK. DH1017.IgM neutralized ZIKV-DAK with a 50% RVP neutralization concentration (RVPNC_50_) of 5.8 ± 2.0 ng/ml (**Figure S6**), which is in line with its ultrapotent neutralization previously described against other ZIKV strains.^30^

To assess the impact of DH1017.IgM on ZIKV IgG-mediated neutralization and vice versa, we measured ZIKV RVP neutralization of DH1017.IgM used at its RVPNC_50_ concentration co-incubated with serially diluted neutralizing and non-neutralizing ZIKV IgG mAbs. Non-ZIKV mAb VRC01, ZIKV IgG mAbs alone and the combination of non-ZIKV-reactive IgM mAb DH1036.IgM ^30^ with ZIKV IgG mAbs served as controls.

Co-incubation of DH1017.IgM with non-neutralizing ZIKV IgG mAbs did not have any adverse effect on DH1017.IgM neutralization, which remained constant at the expected ∼50% regardless of the antibody concentration. (**Figure 2A-D**).

**Figure 2:**
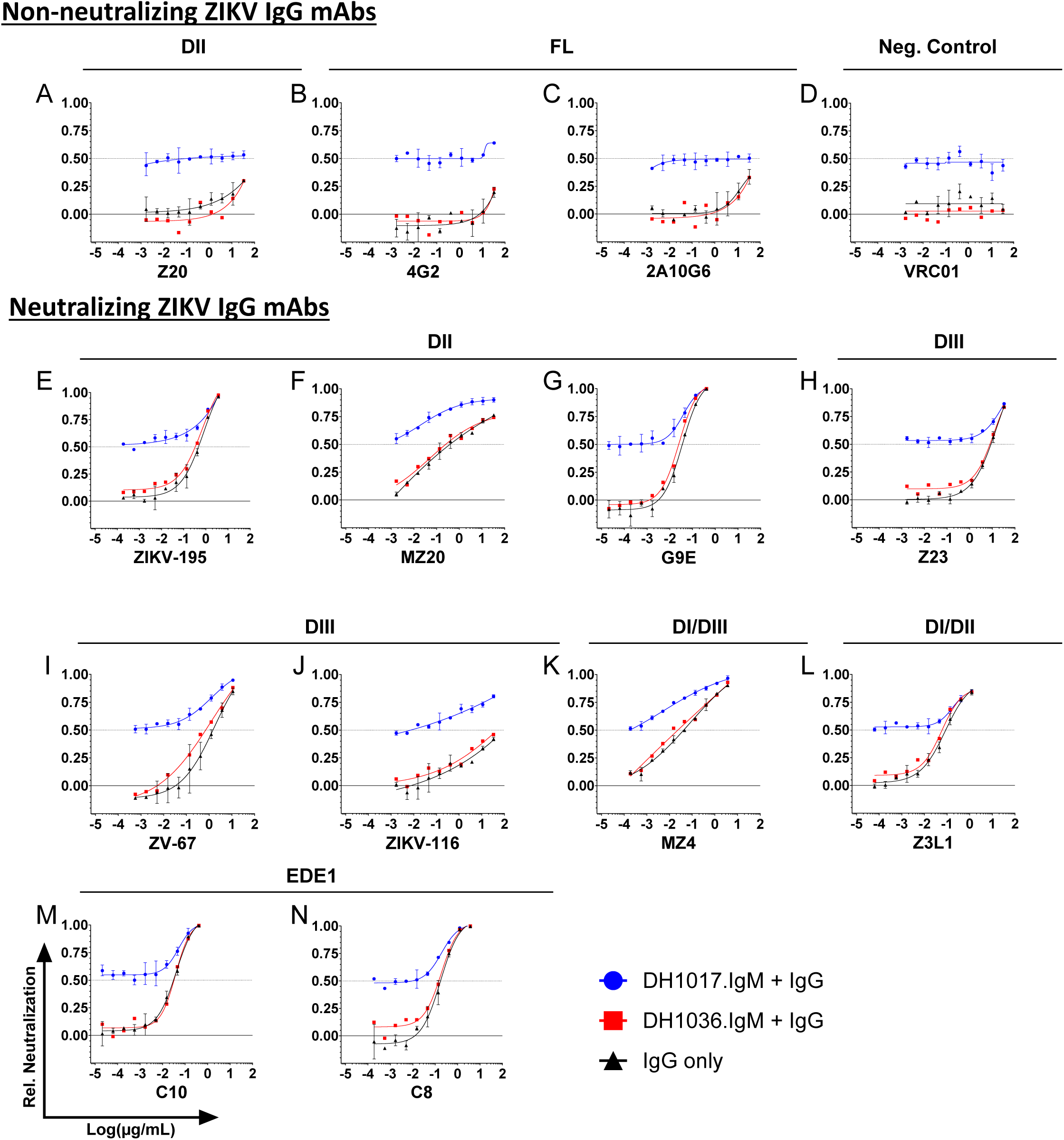
Neutralizing activity of DH1017.IgM in combination with ZIKV-IgG mAbs. Each panel shows the impact of serially diluted (**A-C**) non-neutralizing ZIKV-Ig mAbs, (**D**) non-ZIKV reactive VRC01 mAb, and (**E-N**) ZIKV neutralizing IgG mAbs on the neutralizing activity of a fixed concentration of DH1017.IgM mAb (blue lines) against GFP-expressing ZIKV-DAK reporter virus particle (RVPs) on Raji-DCSIGNR cells. The neutralizing activity of each ZIKV-IgG mAb (black lines) and the combination of ZIKV-IgG mAbs with a non-ZIKV reactive IgM mAbs (DH1036.IgM) are shown in black and red, respectively. The y-axes show neutralizing activity relative to medium-only conditions. Dotted line: 50% neutralization (RVPNC_50_). Error bars: SD of two independent experiments. Panels are sub-grouped by ZIKV-IgG mAb binding regions. See also Figures S4, S5 and S6.

Non-ZIKV-reactive DH1036.IgM did not alter the neutralization profiles of the IgG mAbs when compared to IgG tested alone (**Figure 2**, red and black lines, respectively), whereas combinations of neutralizing ZIKV IgG mAbs with DH1017.IgM displayed at least 50% of viral infectivity neutralization sustained by DH1017.IgM, and improved neutralization tracking the increasing ZIKV-neutralizing IgG concentrations (**Figure 2E-N, blue lines**).

We conclude that DH1017.IgM did not exert obvious detrimental effects on ZIKV IgG-mediated virus neutralization, and vice versa.

### DH1017.IgM reduces ZIKV IgG-mediated ADE activity at sub-neutralizing concentrations

To evaluate if DH1017.IgM affects the ADE activity of ZIKV IgG mAbs, we measured the effect of DH1017.IgM at its fixed RVPNC_50_ concentration on the ADE curve dynamics of each serially diluted IgG (**Figure 3**). Non-neutralizing FL 4G2 and 2A10G6 mAbs, and DII Z20 mAb were poor ADE mediators and coincubation with DH1017.IgM consistently resulted in reduction of IgG-mediated ADE activity (**Figure 3A-C**). For neutralizing ZIKV IgG mAbs, ADE activity was described by a Gaussian curve, which defined a peak ADE activity concentration specific for each mAb (**Figure 3D-M**, black curves). Co-incubation of the neutralizing ZIKV IgG mAbs with non-ZIKV reactive DH1036.IgM did not alter the peak ADE responses (**Figure 3D-M**, red curves). Conversely, co-incubation with DH1017.IgM resulted in a mean 2.7-fold reduction (range: 1.8 – 3.8-fold) in peak ADE infectivity (**Figure 3N**). Importantly, DH1017.IgM did not induce a discernible shift of peak ADE activity for any of the neutralizing ZIKV IgG mAbs toward lower concentrations: rather, a trend toward slightly higher IgG concentrations could be observed (**Figure 3D-M**, blue curves).

**Figure 3:**
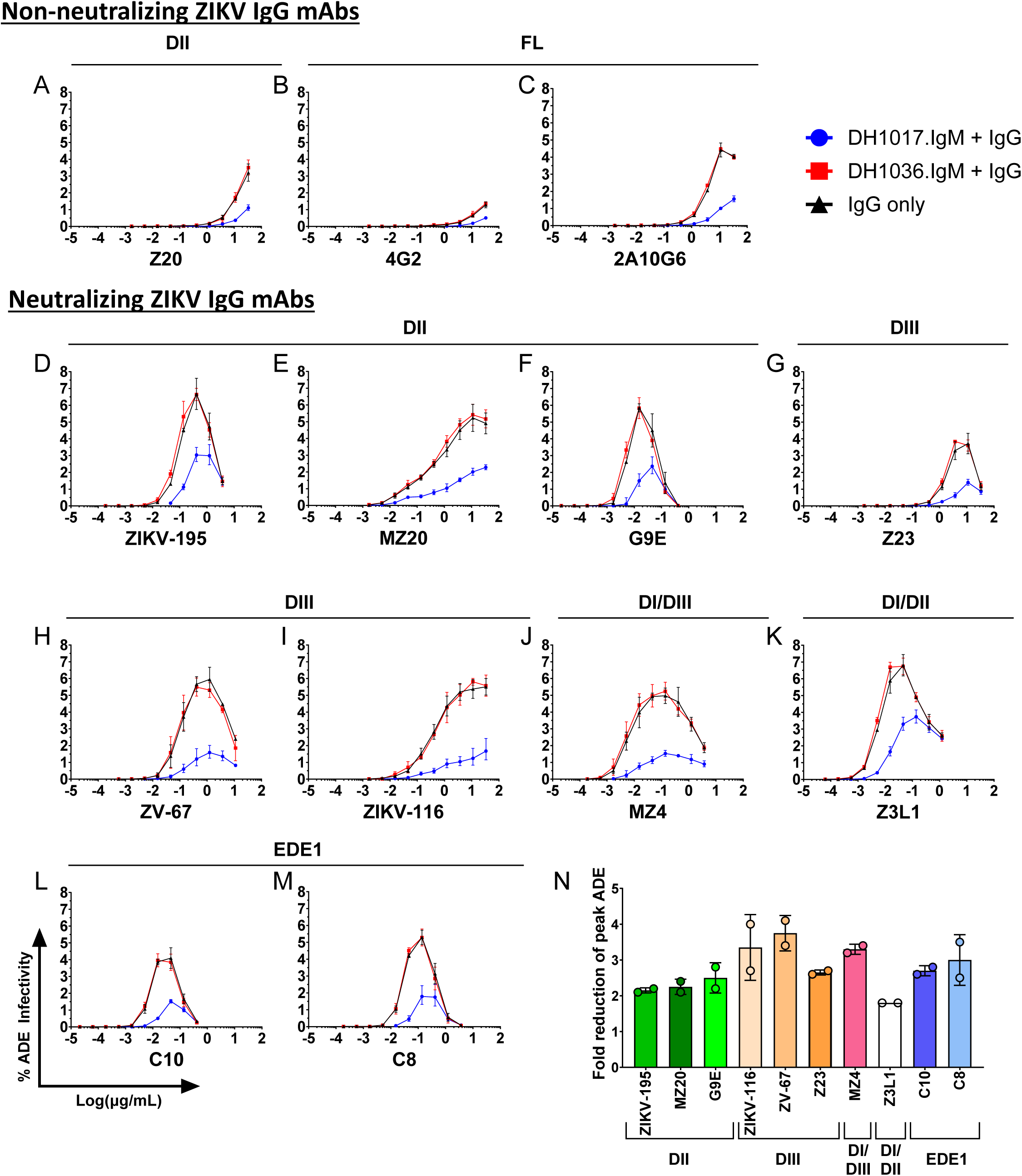
Impact of co-incubation of DH1017.IgM and ZIKV-IgG mAbs on peak antibody-dependent enhancement of infectivity. (**A-M**) Each panel shows ADE-mediated infection of GFP-expressing ZIKV-DAK RVPs by serially diluted (A-C) non-neutralizing and (D-M) neutralizing ZIKV-IgG mAbs either alone (black lines) or in presence of a fixed concentration of DH1017.IgM (blue line) and non-ZIKV reactive DH1036.IgM (red line). The y-axes show the percentage of infected FcγR-expressing K562 over the total number of acquired cells. Error bars show SD of two independent experiments. **(N)** Fold-reduction in ZIKV-IgG peak ADE levels in presence of a fixed concentration of DH1017.IgM. For each ZIKV-IgG, circles show the results of two independent experiments; bars and lines show average and SD. ZIKV-IgG mAbs are grouped and color-coded based on the respective binding regions.

To evaluate DH1017.IgM potency in reducing ZIKV IgG-mediated ADE, we reversed the assay format and co-incubated each IgG mAb at peak ADE concentration (determined as shown in **Figure S4**) with serially diluted DH1017.IgM. Poor ADE mediators 4G2 and Z20 mAbs were excluded from this analysis. DH1017.IgM consistently reduced IgG-mediated ADE activity in a dose-dependent fashion. Remarkably, DH1017.IgM concentrations of 0.1 µg/ml or lower already completely abrogated IgG-mediated ADE, regardless of IgG epitope specificity for all the IgG mAbs in the panel (**Figure 4**). The concentration at which DH1017.IgM reduced 50% of IgG-mediated ADE activity was named cADEC_50_ (where “c” stands for “competing”). Across the IgG panel, DH1017.IgM reduction of IgG-mediated ADE was potent (mean cADEC_50_ = 2.1 ng/ml; range: 0.4 – 9.1 ng/ml). FL 2A10G6 mAb was the least sensitive to DH1017.IgM-mediated ADE reduction (cADEC_50_ = 8.1 ± 1.5 ng/ml) (**Figure 4E**), whereas EDE1 mAbs C8 and C10 were the most sensitive (cADEC_50_ = 0.6 ± 0.4 ng/ml and 0.6 ± 0.3 ng/ml, respectively) (**Figure 4F**).

**Figure 4:**
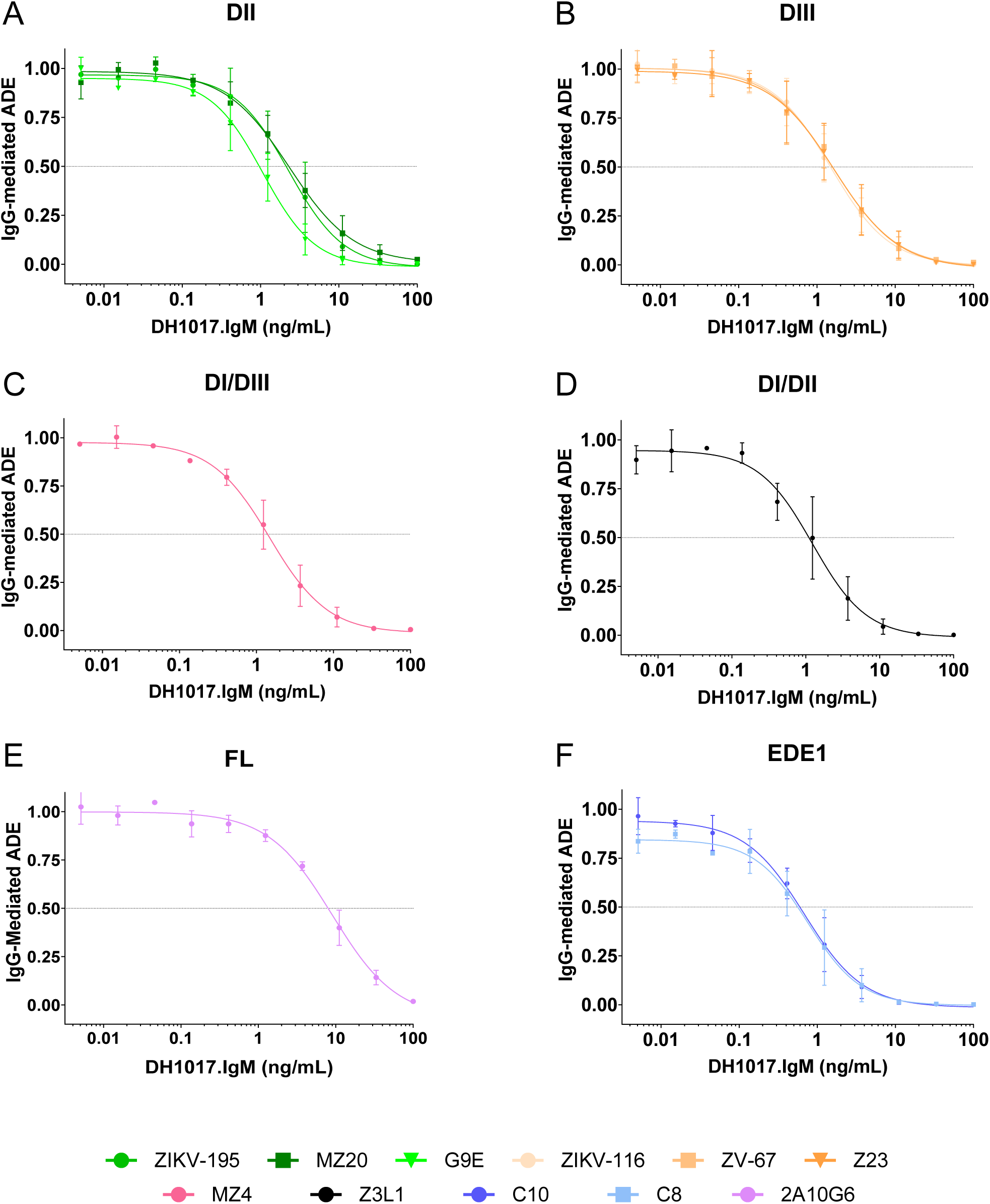
DH1017.IgM dose-dependent potency in reducing ZIKV-IgG mediated ADE infectivity. Each panel shows ADE-mediated infection of GFP-expressing ZIKV-DAK RVPs by a ZIKV-IgG mAb at their respective peak ADE concentration (see Figure S4) in presence of serially diluted DH1017.IgM. The y-axes show the DH1017.IgM dose-dependent reduction in ADE relative to the ZIKV-IgG alone condition. ZIKV-IgG mAbs are grouped in panels based on binding regions: **(A)** DII, **(B)** DIII, **(C)** DI/DIII linker, **(D)** DI/DII hinge, **(E**) fusion loop (FL), and **(F)** EDE1. Error bars: SD from two independent experiments. Dotted line: 50% reduction of ADE-infectivity (cADEC_50_).

In parallel experiments, we measured DH1017.IgM neutralizing activity to compare DH1017.IgM potencies in reducing IgG-mediated ADE (cADEC_50_) and viral neutralization (RVPNC_50_). We defined a conservative threshold of cADEC_50_/RVPNC_50_ ratio ≤ 0.75 to identify reduction of ADE activity at sub-neutralizing concentrations. Remarkably, DH1017.IgM consistently reduced IgG-mediated ADE at sub-neutralizing concentrations with cADEC_50_/RVPNC_50_ ranging from 0.18 for EDE1 C8 and C10 mAbs, to 0.68 for DII MZ20 mAb (**Figure 5A).** DH1017.IgM reduction of FL mAb 2A10G6-mediated ADE narrowly missed the 0.75 threshold and was characterized by a wide variation among replicates (cADEC_50_/RVPNC_50_ = 0.78 ± 0.4) **(Figure 5A**).

**Figure 5:**
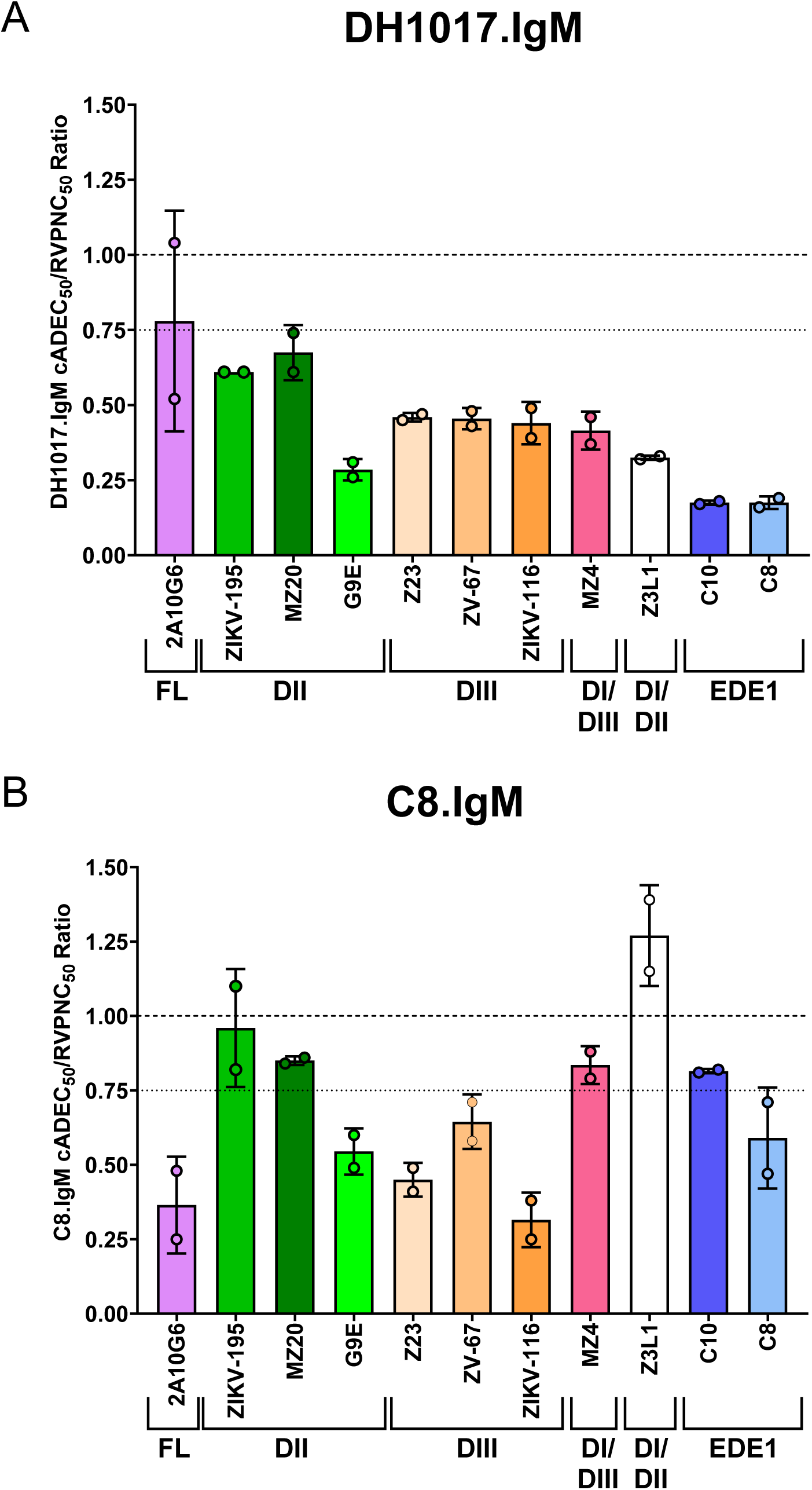
Ratios of ADE mitigation and neutralization potencies of DH1017.IgM and C8.IgM mAbs. The potencies of **(A)** DH1017.IgM and **(B)** C8.IgM to reduce IgG-mediated ADE relative to their respective neutralization capacities were evaluated by calculating the ratio between the cADEC_50_ for each ZIKV-IgG mAb and their RVPNC_50_. The dashed lines indicate a cADEC_50_/RVPNC_50_ ratio of 1 (i.e., no difference). The dotted lines indicate cADEC_50_/RVPNC_50_ = 0.75, which we used to define sub-neutralizing reduction of ADE-mediated infectivity. Error bars: SD from two independent experiments. See also Figures S7 and S8.

We conclude that DH1017.IgM reduces ADE mediated by IgG mAbs recognizing multiple ZIKV E glycoprotein epitopes at sub-neutralizing concentrations.

### IgM reduction of ZIKV IgG-mediated ADE activity is not unique to DH1017.IgM

To verify if the ability to reduce IgG-mediated ADE was exclusive to the DH1017.IgM mAb, we produced and tested a pentameric IgM version on the conformational EDE1 C8 mAb (C8.IgM). C8.IgM neutralized ZIKV-DAK more potently than its IgG counterpart (C8.IgM RVPNC_50_ = 26.3 ± 5.1 pM *vs* C8.IgG RVPNC_50_ = 500 ± 47.3 pM) (**Figure S7**).

C8.IgM successfully mitigated ADE activity of all tested ZIKV-IgG mAbs. As observed for DH1017.IgM, C8.IgM mitigation of IgG-mediated ADE was dose-dependent (**Figure S8**). Similarly to DH1017.IgM, C8.IgM potency in reducing ADE exceeded its neutralization potency in multiple cases (**Figure 5B**). Notably, the profiles of DH1017.IgM and C8.IgM ADE mitigation differed. Like DH1017.IgM, mAb C8.IgM substantially reduced ADE at sub-neutralizing concentrations for 6 of the 11 mAbs, including the autologous C8.IgG, with cADEC_50_/RVPNC_50_ ratios ranging from 0.32 ± 0.09 (ZIKV-116) to 0.65 ± 0.09 (ZV-67) (**Figure 5B**). Conversely, FL 2A10G6-mediated ADE was more potently inhibited by C8.IgM than by DH1017.IgM (cADEC_50_/RVPNC_50_ = 0.37 ± 0.16 vs. 0.78 ± 0.4, respectively), whereas C8.IgM was unable to substantially reduce ADE at sub-neutralizing concentration for DII ZIKV-195 and MZ20, DI/DIII linker region MZ4, EDE1 C10, and DI/DII hinge Z3L1 mAbs. Only Z3L1 mAb displayed an increased cADEC_50_/RVPNC_50_ ratio > 1 (Z3L1 cADEC_50_/RVPNC_50_ = 1.27 ± 0.17) (**Figure 5B**).

Overall, these results demonstrate that the reduction of IgG-mediated ADE by pentameric IgM monoclonal antibodies is a functional property not exclusive to DH1017.IgM.

### DH1017.IgM reduces ADE activity of ZIKV-reactive human sera at sub-neutralizing concentrations

While the polyclonal IgM fraction of sera from ZIKV primary infections can reduce polyclonal IgG-mediated ADE *in vitro*,^32^ it is unknown if a single IgM mAb is sufficient to substantially reduce the polyclonal IgG-mediated ADE. Such distinction is relevant to preliminary assess the potential prophylactic and therapeutic use of monoclonal IgM-derived biologics. Thus, we investigated the ability of DH1017.IgM to reduce ADE activity of unfractionated human sera collected from individuals who participated in a controlled ZIKV infection study (NCT05123222). Study participants were infected with either SJRP/2016-184 (SJRP) (n = 9) or Nicaragua/2016 (NIC) (n = 10) ZIKV strains. Additional seven individuals served as placebo controls. To assess the impact of DH1017.IgM on ADE reduction against a ZIKV strain heterologous to the infecting strain, we used the ZIKV-DAK strain.

At 90 days post-infection, ADE activity for ZIKV-DAK strain was measurable in all ZIKV-SJRP and ZIKV-NIC infected sera with peak titers between 1,215 and 3,645, respectively, and absent in placebo controls (**Figure 6A,B**). Sera at peak ADE titers were co-incubated with serially diluted DH1017.IgM to measure its effect on serum ADE activity. Similarly to what was observed with ZIKV IgG mAbs, DH1017.IgM potently reduced serum-mediated ADE in a dose-dependent manner, with mean DH1017.IgM cADEC_50_ of 6.2 ± 2.2 ng/ml in the ZIKV-SJRP cohort (**Figure 6C**) and 6.0 ± 2.6 ng/ml in the ZIKV-NIC cohort (**Figure 6D**). The DH1017.IgM cADEC_50_/RVPNC_50_ ratios were 0.62 ± 0.2 for the ZIKV-SJRP cohort and 0.60 ± 0.3 for the ZIKV-NIC cohort, with no statistically significant difference between the two groups (p=0.8821, unpaired Student’s t-test) (**Figure 6E**). Remarkably, DH1017.IgM reduced serum-mediated ADE at sub-neutralizing concentrations (cADEC_50_/RVPNC_50_ ≤ 0.75) in 84.2% (16 out of 19) of the tested sera (**Figure 6E**).

**Figure 6:**
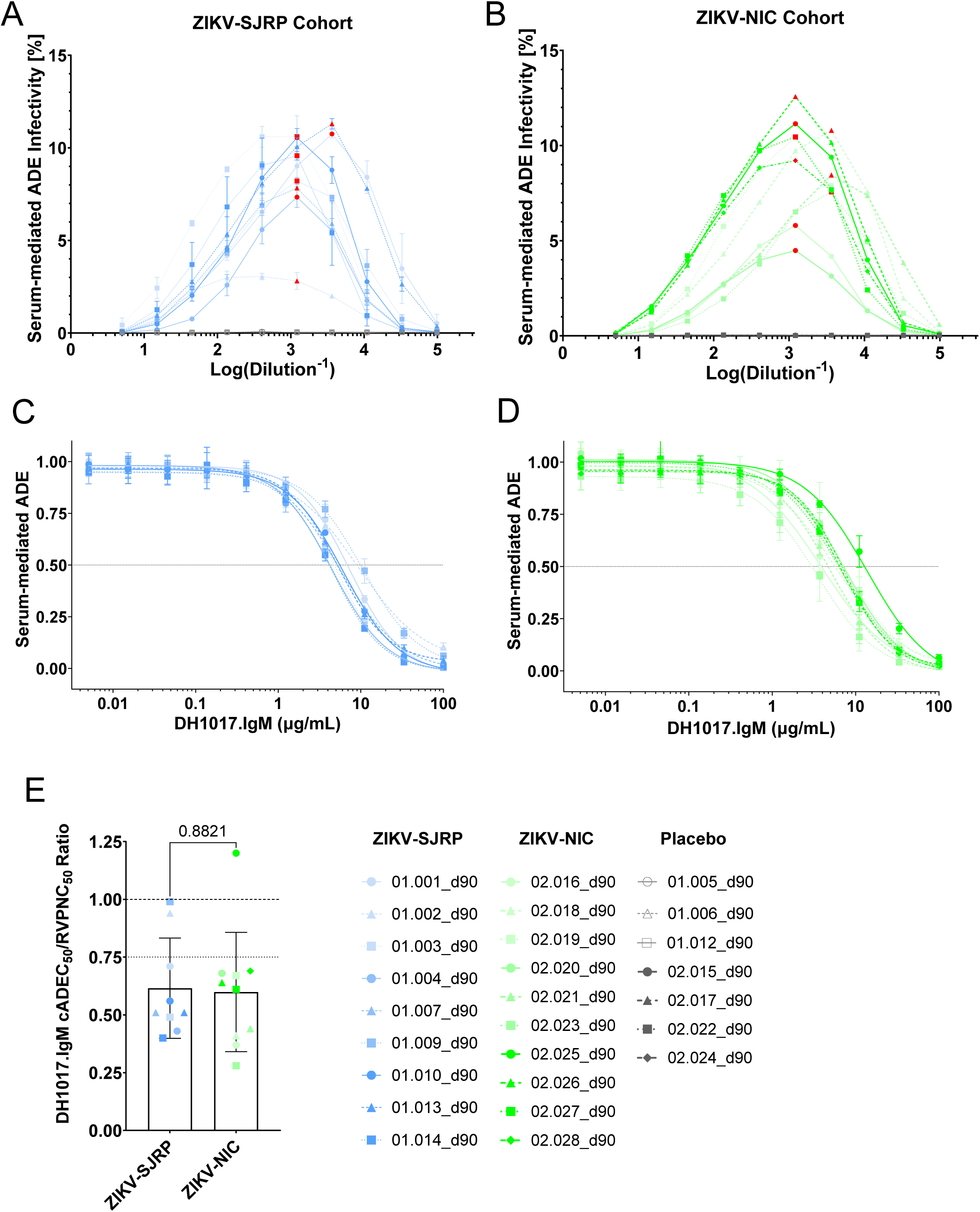
DH1017.IgM reduces ADE infectivity mediated by ZIKV polyclonal human sera. The ADE-mediated infection of GFP-expressing ZIKV-DAK RVPs for human sera collected 90 days post inoculation with **(A)** ZIKV-SJRP or **(B)** ZIKV-NIC are reported as percentage of infected FcγR-expressing K562 cells over the total number of acquired cells. Red data points identify the ADE-peak titer for each serum or, where no clear peak titer could be identified, the highest dilution with maximum ADE infectivity, which was used in subsequent ADE-competition assays. **(C,D)** ADE-mediated infection of GFP-expressing ZIKV-DAK RVPs by (C) ZIKV-SJRP- and (D) ZIKV-NIC infected sera at their peak ADE titers in presence of serially diluted DH1017.IgM. The y-axes show the DH1017.IgM dose-dependent reduction in ADE relative to the serum-only condition. Error bars: SD of two independent experiments. Dotted line: 50% reduction of ADE-infectivity (cADEC_50_). **(E)** DH1017.IgM cADEC_50_/RVPNC_50_ ratio for each serum. Symbols: Average ratio of two independent experiments; bar: group mean; error bars: SD of all sera within each group. Statistical significance was evaluated using unpaired Student’s t-test.

We conclude that DH1017.IgM reduced ADE activity of human sera from ZIKV-infected individuals, typically at sub-neutralizing concentrations.

## DISCUSSION

ZIKV infections are characterized by prolonged serum IgM titers. We and others have described that serum ZIKV IgM contributes to ZIKV neutralization; moreover, serum Ig-depletion studies implicated primary IgM responses to ZIKV infection in mitigating IgG-mediated ADE. However, the manner in which ZIKV IgM exerts its functions in the presence of ZIKV IgG mAbs remained incompletely defined. We previously described ZIKV mAb DH1017.IgM, a prototypic ZIKV-specific IgM in its native multimeric conformation, whose ultrapotent neutralization was supported by multimeric modes of antigen recognition exclusive to the IgM isotype. In this study, we investigated how co-incubation of DH1017.IgM and ZIKV IgG mAbs that target multiple ZIKV E-dimer regions reciprocally alters virion binding, viral neutralization, and ADE activities. We demonstrated that antibodies of the IgM isotype reduce IgG-mediated ADE regardless of ZIKV epitope specificity, in a dose-dependent manner and at sub-neutralizing concentrations, hence identifying IgG-mediated ADE reduction as an independent protective IgM function that is distinct from viral neutralization and likely multifactorial.

We had previously speculated that, while the greater avidity of DH1017.IgM could outcompete less avid IgG antibodies, the large footprint associated with its multivalent mode of virion recognition could also represent a disadvantage in the presence of ZIKV IgG antibodies.^30^ Here, we describe a complex cross-competition dynamic between DH1017.IgM and ZIKV IgG mAbs: IgG mAbs targeting all domain specificities competed, at least partially with DH1017.IgM binding (i.e., DIII mAbs did not reach 50% DH1017.IgM blocking). However, based on the respective IC_50_ concentrations, DH1017.IgM outcompeted most ZIKV IgG mAbs, predicting a potential advantage of DH1017.IgM in a polyclonal environment. That DH1017.IgM successfully abrogated IgG-mediated ADE activity in sera of ZIKV-infected individuals further corroborates this hypothesis. Nonetheless, at saturating concentration, DH1017.IgM reduced, but did not abrogate, IgG virion binding in competition assays. This finding implies that the ability of DH1017.IgM to compete ZIKV IgG virion binding is self-limited and it is compatible with the previously proposed theoretical maximum of three DH1017.IgM mAbs concurrently binding to 30 of the 90 epitopes on the same virion and that such an arrangement could leave epitopes still accessible to smaller IgG mAbs across the virion surface.^30^ Importantly, the cross-competition between DH1017.IgM and ZIKV IgG mAbs did not exert negative effects on viral neutralization.

The incomplete blocking of ZIKV IgG mAbs implied that DH1017.IgM affects the stoichiometry of IgG binding to ZIKV, which is a critical factor for ADE of flaviviruses.^33^ Indeed, DH1017.IgM reduced IgG-mediated ADE independent of the IgG epitope specificity. Co-incubation of ZIKV IgG mAbs and DH1017.IgM resulted in IgG-mediated ADE peak activities shifted, albeit modestly, toward higher IgG concentrations. We speculate that this phenomenon could contribute to IgM-mediated mitigation of ADE *in vivo* when ADE is sustained by waning IgG immunity. Due to the ample cross-reactivity of humoral responses elicited by ZIKV and DENV infections, and since many of the IgG mAbs included in the panel were elicited by and/or cross-reacted with DENV, the same protective function could be at play when cross-reactive IgG antibodies are recalled during sequential heterologous infections.

Notably, for DI/DIII MZ4, DIII ZIKV-116 and ZV-67 IgG mAbs, which were not efficiently competed by IgM, the shift in IgG-mediated ADE peak concentration was not observed, and yet, they displayed the strongest ADE peak reduction. This finding suggests that IgM-mediated mitigation of ADE is multifactorial and may involve additional yet-to-be-defined mechanisms.

Previous studies on fractionated sera of ZIKV infected individuals implicated ADE reduction by polyclonal IgM.^32^ Here, we demonstrate that IgM mAb can potently inhibit ADE activity. Critically, DH1017.IgM retained the ability to reduce ADE even at sub-neutralizing concentrations in most of the sera obtained from ZIKV-infected individuals, indicating that the presence of polyclonal serum IgM had negligible effects on its ADE inhibiting activity. For the few exceptions, we cannot exclude that serum IgM already exerted some ADE-mitigating activity, hence marginally reducing the impact of DH1017.IgM. That a recombinantly produced IgM mAb, EDE1 C8.IgM, also broadly reduced IgG-mediated ADE demonstrates that this is not a function exclusive to DH1017.IgM. Furthermore, differences in the ADE mitigation profiles between DH1017.IgM and C8.IgM imply different interaction dynamics between IgG and IgM pairs, which can be complimentary, as most dramatically evidenced by the divergent DH1017.IgM and C8.IgM cADEC_50_/RVPNC_50_ ratios for mAbs 2A10G6 and Z3L1.

These observations provide the rationale to consider multivalent IgM-based biologics as countermeasures to protect from the severe effects of flavivirus-mediated ADE *in vivo*. In Zika, ADE may play a role in supporting vertical transmission.^8,10–12^ Combinations of potently neutralizing ZIKV IgG mAbs did not fully protect fetuses from vertical transmission in non-human primate studies.^34,35^ Given the combined functions of IgM for potent ZIKV neutralization, potent inhibition of ADE activity, and the inability to mediate virus infectivity of monocytes and macrophages or transport opsonized viruses across the placenta, future studies should explore the suitability of IgM-derived biologics to prevent ZIKV vertical transmission.

In addition, since the virion architecture and E-protein structure of ZIKV and DENV are highly similar, and ADE is believed to contribute to the development of severe dengue upon secondary DENV infections, this study provides a rationale to investigate IgM antibody-based ADE mitigation strategies aimed at treating or preventing severe dengue. Since DH1017.IgM does not bind to DENV,^30^ we could not test its functions in the context of dengue infections. Future studies should investigate if DENV-reactive IgM mAbs can also efficiently mitigate IgG-mediated ADE of DENV infectivity across multiple serotypes.

A limitation of this study is that we performed *in vitro* and *ex vivo* experiments but not *in vivo* studies. The interpretation of *in vivo* inhibition of ADE activity will likely be complicated by DH1017.IgM ultrapotent neutralization and it should be addressed in future studies.

In conclusion, this study describes the complex functional interactions between ZIKV IgM and IgG antibodies at monoclonal level, demonstrates that ZIKV mAbs of the IgM isotype reduce IgG-mediated ADE regardless of ZIKV epitope specificities, that such protective function is distinct from viral neutralization and likely multifactorial, and suggests that multivalent IgM-based biologics may offer a promising countermeasure against ZIKV infection and vertical transmission.

## METHODS

### Cell lines

The K562-FcγR and Raji DC-SIGNR cell lines were kindly provided by Dr. Theodore C. Pierson (VRC, NIAID). Cells were maintained in RPMI medium with 1x Glutamax (Gibco 61870-036) supplemented with 10% (v/v) heat-inactivated fetal bovine serum (FBS) (ThermoFisher, Gibco A5669801). Culture flasks were incubated in a humidified incubator at 37°C, 5% CO_2_. All assays were performed in RPMI media with 1x GlutaMax without phenol-red (Gibco 11835-030) supplemented with 10% (v/v) FBS.

### Monoclonal antibody production

All recombinant IgG mAbs were expressed using heavy chain expression plasmid pFUSEss-CHIg-hG1 (InvivoGen) and the respective light chain expression plasmid pFUSE2ss-CLIg-hL2 (InvivoGen) or pFUSE2ss-CLIg-hk (InvivoGen) for lamba- and kappa-light chains, respectively, and transfected into Expi293F cells (ThermoFisher) with the Gibco Expi293F Expression Kit (ThermoFisher) following manufacturer’s protocol. Cell supernatants were harvested 5-7 days post transfection, clarified by centrifugation, and mAbs were purified with Protein G columns (Cytivia) either by gravity flow or on the AktaGo FPLC instrument (Cytivia). Eluates were buffer exchanged to PBS using Amicon Ultra-15 Centrifugal Filters, Ultracel, 50k MWCO (Millipore). Purified mAbs were quantified on a Cytation 7 using a Take3 plate (Agilent) and stored at -80°C until use. DH1017.IgM and DH1036.IgM mAbs were expressed by B lymphoblastoid cell lines.^30^ IgM mAbs were purified using POROS CaptureSelectIgM-XL Affinity Matrix (Invitrogen) and the eluates were buffered exchanged using Vivaspin 20, 300k MWCO PES (Sartorius). Purified IgM mAbs were quantified on a Cytation 7 using a Take3 plate (Agilent) and stored at 4°C until use. Monoclonal antibody C8.IgM was commercially produced by SinoBiological. Each antibody lot was validated through mass photometry (MP) and whole virion binding ELISA.

### Conjugation of ZIKV-reactive monoclonal antibodies to horse radish peroxidase (HRP)

HRP-conjugation was performed by labeling each mAb with HRP Conjugation Kit-Lightning-Link (ab102890) according to manufacturer’s instructions. Briefly, mAbs were diluted with sterile 1x PBS at the concentration of 1 mg/mL. For each 10 µL of mAbs, 1 µL of Modifier reagent was added, and then the sample was used to resuspend the lyophilized HRP. HRP-conjugation was allowed to proceed for 4 h in the dark at room temperature. Then, reaction was quenched by adding 1 µL of Quencher reagent for every 10 µL of mAbs. HRP-conjugated mAbs were stored at 4°C overnight, before being 0.22 µm-sterile filtered. Each lot was titrated to confirm binding to ZIKV whole virions.

### Whole Virion Binding ELISA

High-binding 384-well microplates (Corning, #3700) were coated overnight at 4°C with 15 ng/well of 4G2 mAb (Curia) in 0.1 M NaHCO_3_ buffer. Plates were blocked in assay diluent [1x PBS, 15% (v/v) normal goat serum, 2% (w/v) whey protein (Sigma, W15000), 0.5% (v/v) Tween-20, and 0.05% (v/v) NaN_3_] for 1 h at 37°C, followed by an incubation with a previously optimized dilution of ZIKV/Aedes-africanus/SEN/DakAr41524/1984 (ZIKV-DAK) for 1 h at 37°C. Primary purified mAbs (10 µL/well) were added and incubated for 1 h at 37°C. Each mAb was tested in duplicate using ten-points of a 4-fold serial dilution in assay diluent, starting at 100 µg/mL. Horseradish peroxidase (HRP)-conjugated goat anti-human IgG (Jackson ImmunoResearch Laboratories, #109-035-098, 1:6000), anti-human IgA (Jackson ImmunoResearch Laboratories, #109-035-011, 1:6000), or anti-human IgM (Jackson ImmunoResearch Laboratories, #109-035-129, 1:10,000) secondary antibodies were used in assay diluent without NaN_3_ and incubated for 1 h at 37°C, followed by the addition of SureBlue Reserve TMB substrate (KPL, #5120-0083). Each secondary antibody was used at previously optimized lot-specific concentrations. Reactions were stopped using 0.1 M HCl. Optical density (OD) at the wavelengths of 450 nm and 650 nm -for antibody quantification and detection of aspecific OD spikes, respectively -were measured on a BioTek Cytation 7 microplate reader (Agilent). All wash steps were performed with 1x PBS, containing 0.1% Tween-20 using the BioTek EL406 Washer Dispenser (Agilent) and all reagents were added using either the Biomek i7 automated liquid handling system (Beckman Coulter), Voyager multichannel pipette (Integra), or the 384-channel ViaFlo384 electronic pipetting system (Integra). OD_450_ values were used to calculate EC_50_ titers through a four-parameter nonlinear regression analysis with variable slope in Prism 10 (GraphPad).

### Whole Virion Binding Competition ELISA

ELISA-based binding competition assays were performed as described for Whole Virion Binding ELISA with the following modifications: Primary unconjugated mAbs were brought to 100 µg/mL in assay diluent, further diluted 4-fold for 9 steps and added to the plates (10 µL/well) in technical duplicates for 1 h incubation at 37°C. ZIKV-reactive HRP-mAbs were diluted in assay diluent without NaN_3_ at their pre-determined lot-specific whole virion binding EC_50_. Upon one single washing step, each diluted ZIKV-reactive HRP-mAb was added to the plate (10 µL/well) and incubated for 1 h at 37°C. Negative controls of binding competition included assay diluent alone and non-ZIKV-reactive HRP-mAbs VRC01 or DH1036.IgM. Autologous blocking mediated by each primary unconjugated mAb against its own HRP-conjugated version was used as positive control of binding competition.

Upon background subtraction, binding competition values were calculated as percentages using the following formula: (1 -[mean OD_450nm_ (Unconjugated mAb dilution) – mean OD_450nm_ (HRP-conjugated mAb only)] x 100). Binding competition values were used to perform a four-parameter nonlinear regression analysis with variable slope in Prism 10 (GraphPad) and the absolute 50% inhibitory concentration (IC_50_) was calculated.

### RVP-based fluorescent neutralization assays

MAb neutralization was determined using a flow cytometry-based neutralization assay in Raji cells expressing DC-SIGNR with reporter virus particles (RVPs), as previous described ^36,37^. We produced RVPs containing GFP reporter gene and expressing the structural proteins (CprME) of ZIKV/Aedes-africanus/SEN/DakAr41524/1984 (ZIKV-DAK) (GenBank: KX601166.2). Serial three-fold dilutions of mAbs were mixed with an equal volume of RVPs diluted in assay medium (RPMI medium without phenol red, supplemented with 10% FBS and 1x Anti-Anti) to achieve 2% -5% infection of total cells and incubated at 37°C. After 1h, 5×10^5^ cells/ml of Raji DC-SIGNR cells were added to each mAb-RVP mixture in an equal volume of assay medium and incubated for 48 h at 37°C, in 5% CO_2_, humidified atmosphere. Neutralization assays were performed in a total volume of 150µLs or 300µLs. The percentage of infected cells were quantified on a BD FACSLyric, BD Symphony, or BD FACS CantoII (BD Biosciences).

Data were analyzed in FlowJo v10.9.0 (BD Biosciences) and titration curves were fitted using a 4-point logistic regression model in Prism 10 (GraphPad). The 50% RVP neutralization concentration (RVPNC_50_) was defined as the mAb concentration that reduced infectivity by 50% compared to the medium only control condition. All assays were performed in two independent replicates and the mean RVPNC_50_ and corresponding standard deviation is reported.

To measure the combinatorial effect of DH1017.IgM and individual IgG mAbs, DH1017.IgM or DH1036.IgM (negative control) were diluted to twice the concentration of the of DH1017.IgM RVPNC_50_. Serial three-fold dilutions of IgG mAbs were added at equal volumes to the IgM dilution, resulting in a constant concentration of DH1017.IgM at its RVPNC_50_ and a decreasing concentration of IgG.

### RVP-based fluorescent ADE assays

*In vitro* antibody-dependent enhancement (ADE) of ZIKV infection was quantified using flow cytometry-based RVP assay with K562-Fcγ-RII cells, as previously described ^36^. The ADE assay setup was identical to the RVP-based neutralization assays described above, including the strategy to measure the combinatorial effect of a fixed amount of DH1017.IgM on each IgG mAb ADE activity, except for the cell type used.

To measure the dose-dependent effect of DH1017.IgM on IgG-mediated ADE, IgG mAbs were brought to twice the peak ADE concentrations and were combined with an equal volume of serially diluted DH1017.IgM starting at 0.1 µg/mL (three-fold serial dilutions, 10 steps), resulting in a constant IgG ADE peak concentration and a decreasing concentration of DH1017.IgM. ADE competition assays using serially diluted DH1017.IgM were performed in a total volume of 150µL. Data acquisition and analysis were performed as reported above. The concentration of DH1017.IgM that reduced by 50% the ADE activity of each ADE-mediating mAbs alone was defined cADEC_50_ (were “c” stands for “competing”).

### Human serum from ZIKV control human infection model study CIR 316

From the controlled human infection model study CIR 316 (NCT05123222), we obtained 18 sera collected from female subjects 90 days post subcutaneous inoculation of placebo (n=7), 100 PFUs of ZIKV-SJRP/2016-184 (n=9), or 100 PFUs of ZIKV-Nicaragua/2016 (n=10). Serum samples were heat-inactivated by incubation at 56°C for 30 min prior to use in the assays described above. The clinical protocol was approved by the Johns Hopkins School of Public Institutional Review Board and all volunteers signed informed consent documents.

## RESOURCE AVAILABILITY

### Lead contact

Further information and requests for resources and reagents should be directed to and will be fulfilled by the lead contact, Mattia Bonsignori.

### Materials availability

The research materials used in this study are available, contingent to product availability, upon written request and subsequent execution of an appropriate materials transfer agreement.

### Data and code availability

Sequences of Ig V(D)J rearrangements of all antibodies used in this study are publicly available in GenBank. Accession numbers are listed in the key resource table. This paper does not report original antibody sequences or code. Any additional information required to reanalyze the data reported in this paper is available from the lead contact upon request.

## ACKNOWLEDGEMENTS

The authors thank the Dr. Grzegorz Piszcek and Dr. Di Wu from the Biophysics Core Facility at National Heart, Lung, and Blood Institute (NHLBI) of the National Institutes of Health (NIH) for their advice and expertise on mass photometry and for making the mass photometry instrument available for use; and Dr. Anna Durbin (Center for Immunization Research, Johns Hopkins University) for providing human sera from the controlled human infection model study CIR 316 (NCT05123222).

This research was supported by the Intramural Research Program of the National Institutes of Health (NIH). The contributions of the NIH authors are considered Works of the United States Government. The findings and conclusions presented in this paper are those of the authors and do not necessarily reflect the views of the NIH or the U.S. Department of Health and Human Services.

## Author Contributions

Conceptualization: M.L., M.B.; Data curation, M.L., T.J.M., G.S.M.; Formal analysis: M.L., T.J.M, G.S.M., M.B; Funding acquisition: M.B.; Investigation: M.L., T.J.M., G.S.M.; Methodology: M.L., T.J.M., G.S.M.; M.B.; Project administration: M.L., M.B.; Resources: S.S.W., J.M.D., M.B.; Supervision: M.L.,M.B.; Validation: M.L., M.B.; Visualization: M.L., T.J.M., G.S.M., M.B.; Writing – original draft: M.L., T.J.M., G.S.M, M.B; Writing – review & editing: M.L., T.J.M, G.S.M., J.M.D., S.S.W., M.B.

## Supplemental information titles and legends

Figures S1-S8

## Key resources table

| REAGENT or RESOURCE | SOURCE | IDENTIFIER |
| --- | --- | --- |
| Antibodies |  |  |
| DH1017.IgM | Produced from B-LCL in-house | Singh et al. 2022<sup>30</sup> |
| DH1036.IgM | Produced from B-LCL in-house | Singh et al. 2022<sup>30</sup> |
| C8.IgM | Custom production by Sino Biological | N/A |
| Horseradish peroxidase (HRP)-conjugated goat anti-human IgG | Jackson ImmunoResearch Laboratories | Cat# 109-035-098 |
| HRP-conjugated goat anti-human IgM secondary antibody | Jackson ImmunoResearch Laboratories | Cat# 109-035-129 |
| VRC01 | Kindly provided by Dr. Tongqing Zhou, VRC/NIAID/NIH | GenBank: GU980702, GU980703 |
| Z3L1.IgG | Recombinantly produced in this study | N/A |
| G9E.IgG | Recombinantly produced in this study | N/A |
| ZIKV-195.IgG | Recombinantly produced in this study | N/A |
| MZ20.IgG | Recombinantly produced in this study | N/A |
| Z20.IgG | Recombinantly produced in this study | N/A |
| 2A10G6.IgG | Recombinantly produced in this study | N/A |
| Z23.IgG | Recombinantly produced in this study | N/A |
| ZV-67.IgG | Recombinantly produced in this study | N/A |
| ZIKV-116.IgG | Recombinantly produced in this study | N/A |
| ZV-64.IgG | Recombinantly produced in this study | N/A |
| MZ4.IgG | Recombinantly produced in this study | N/A |
| C8.IgG | Recombinantly produced in this study | N/A |
| C10.IgG | Recombinantly produced in this study | N/A |
| 4G2.IgG | Produced by Curia - |  |
| DH1017.IgG | Recombinantly produced in this study | N/A |
| DH1017.IgM | Recombinantly produced in this study | N/A |
| Bacterial and virus strains |  |  |
| ZIKV/Aedes-africanus/SEN/DakAr41524/1984 (ZIKV-DAK) | Kindly provided by Dr. Matthew Aliota, University of Minnesota |  |
| Biological samples |  |  |
| Human serum from controlled human infection model study CIR 316 | Kindly provided by Dr. Stephen S. Whitehead, LVD/NIAID/NIH | NCT05123222 |
| Chemicals, peptides, and recombinant proteins |  |  |
| Whey protein | Sigma | Cat# W15000 |
| Normal goat serum | Thermo Fisher | Cat# 16210-072 |
| POROS™ CaptureSelect™ IgM-XL Affinity Matrix | Invitrogen | Cat# 2812892050 |
| HiTrap Protein G HP antibody purification columns, 1 ml | Cytivia | Cat# 29048581 |
| Critical commercial assays |  |  |
| HRP Conjugation Kit-Lightning-Link | Abcam | Cat# ab102890 |
| Deposited data |  |  |
| Z3L1 | GenBank | KX929926.1, KX929927.1 |
| G9E | PDB | 7YAR |
| ZIKV-195 | PDB | 6MID |
| MZ20 | GenBank | MN523670.1,<br>MN523677.1 |
| Z20 | GenBank | KX929924.1,<br>KX929925.1 |
| 4G2 | GenBank | KJ438785.1,<br>KJ438784.1 |
| 2A10G6 | PDB | 5JHL |
| Z23 | GenBank | KX929928.1,<br>KX929929.1 |
| ZV-67 | PDB | 5KVG |
| ZIKV-116 | PDB | 6PLK |
| ZV-64 | PDB | 5KVF |
| MZ4 | GenBank | MN523667.1,<br>MN523674.1 |
| C8 | PDB | 4UTA |
| C10 | PDB | 5H37 |
| DH1017 | GenBank | OP612789,<br>OP612798 |
| CprME protein sequence of ZIKV/Aedes-africanus/SEN/DakAr41524/1984 (ZIKV-DAK) | GenBank | KX601166.2 |
| Experimental models: Cell lines |  |  |
| K-562 cells | Kindly provided by<br>Dr. Theodore C.<br>Pierson,<br>VRC/NIAID/NIH | ATCC CCL-243 |
| Raji DC-SIGNR | Kindly provided by<br>Dr. Theodore C.<br>Pierson,<br>VRC/NIAID/NIH | Davis et al., 2006 <sup>46</sup> |
| Vero African green monkey kidney epithelial cells | ATCC | ATCC CCL-81 |
| DH1017.IgM B-LCL | Singh et al., 2022 <sup>30</sup> | N/A |
| DH1036.IgM B-LCL | Singh et al., 2022 <sup>30</sup> | N/A |
| Recombinant DNA |  |  |
| pFUSEss-CHlg-hG1 | InvivoGen | Cat# pfusess-hchg1 |
| pFUSE2ss-CLlg-hL2 | InvivoGen | Cat# pfuse2ss-hcll2 |
| pFUSE2ss-CLlg-hk | InvivoGen | Cat# pfuse2ss-hclk |
| Software and algorithms |  |  |
| FlowJo | BD Biosciences | v10.9.0 |
| GraphPad Prism 10 | GraphPad Software | v10 |
| BD FACSDiva Software | BD Biosciences | v9.0 |

