## Supplemental Figures for "Sub-neutralizing concentrations of Zika virus IgM monoclonal antibodies reduce IgG-mediated antibody-dependent enhancement of infectivity"

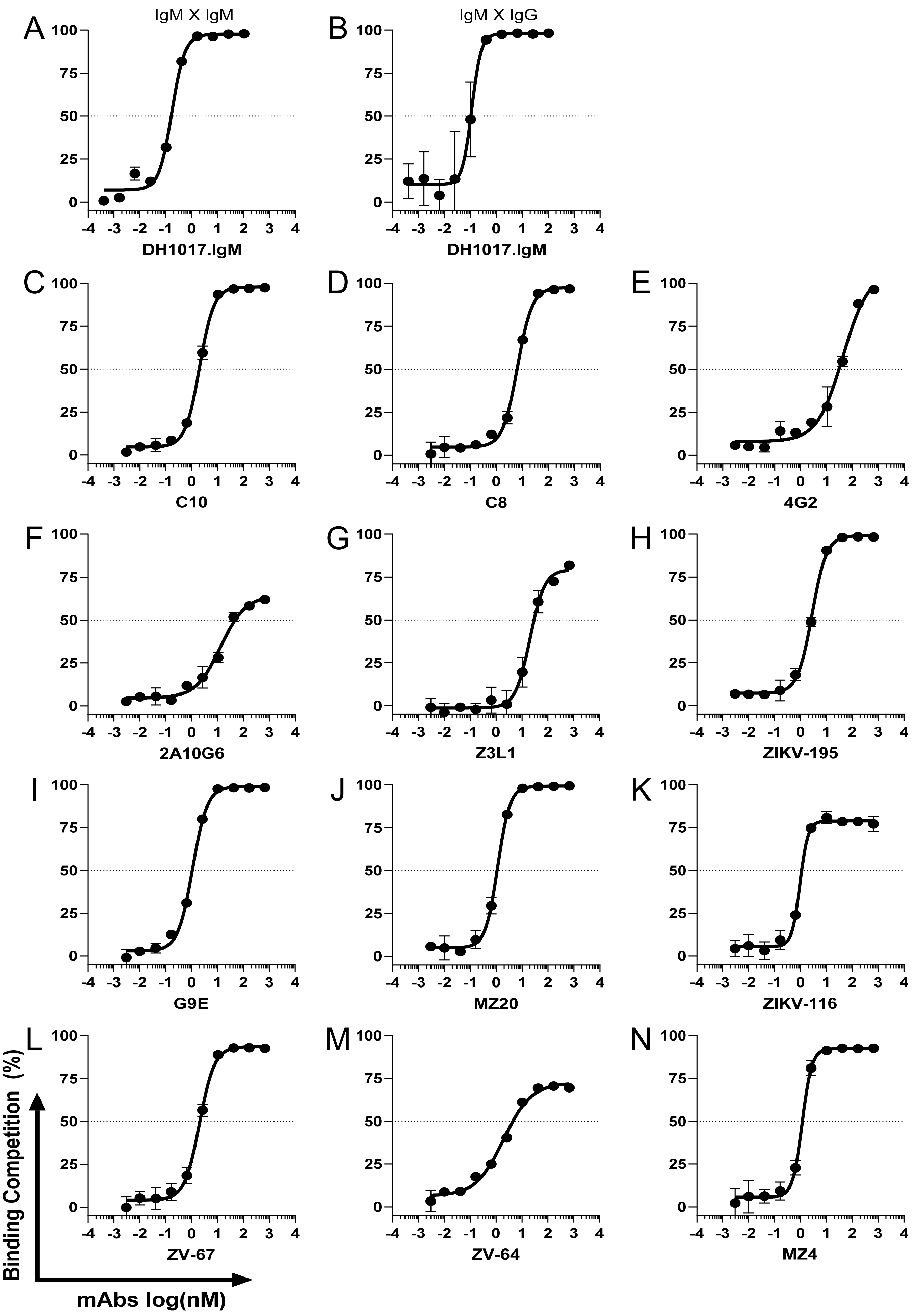

A
IgM X IgM
DH1017.IgM
B
IgM X IgG
DH1017.IgM
C
C10
D
C8
E
4G2
F
2A10G6
G
Z3L1
H
ZIKV-195
I
G9E
J
MZ20
K
ZIKV-116
L
ZV-67
M
ZV-64
N
MZ4
Binding Competition (%)
mAbs log(nM)

**Figure S1. ZIKV-DAK whole virion binding autologous competition of DH1017.IgM and ZIKV-IgG mAbs.** (**A and B)** Validation of the ability of unconjugated DH1017.IgM to compete binding to ZIKV-DAK whole virions by its own HRP-conjugated versions expressed as an IgM (A) or as an IgG (B). **(C-N)**: Validation of the ability of each unconjugated ZIKV-IgG mAbs to compete binding of their own HRP-conjugated versions. Dotted lines: 50% binding reduction of HRP-conjugated mAbs alone. Error bars: SD of two technical replicates. Data are representative of at least two independent experiments.

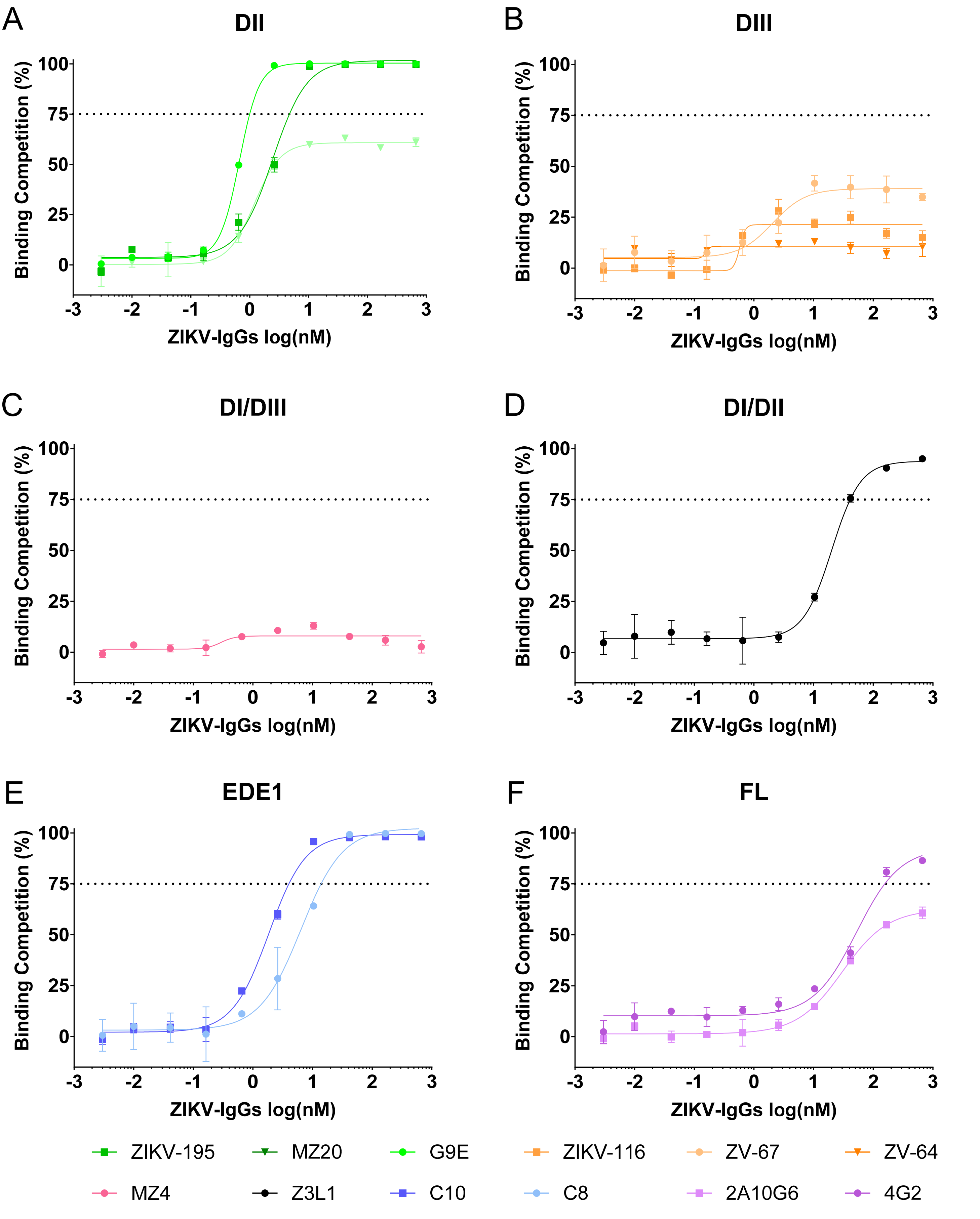


**Figure S2. ZIKV-IgG dose-dependent competition of DH1017.IgM binding to ZIKV-DAK whole virion.** Dose-response curves show the percentage of DH1017.IgM binding to ZIKV-DAK competed by each of the 12 ZIKV-IgG mAbs grouped in panels based on binding regions: **(A)** DII, **(B)** DIII, **(C)** DI/DIII linker, **(D)** DI/DII hinge, **(E)** EDE1, and **(F)** fusion loop (FL). Dotted lines: 75% binding reduction of DH1017.IgM binding. Error bars: SD of two technical replicates. Data are representative of two independent experiments.

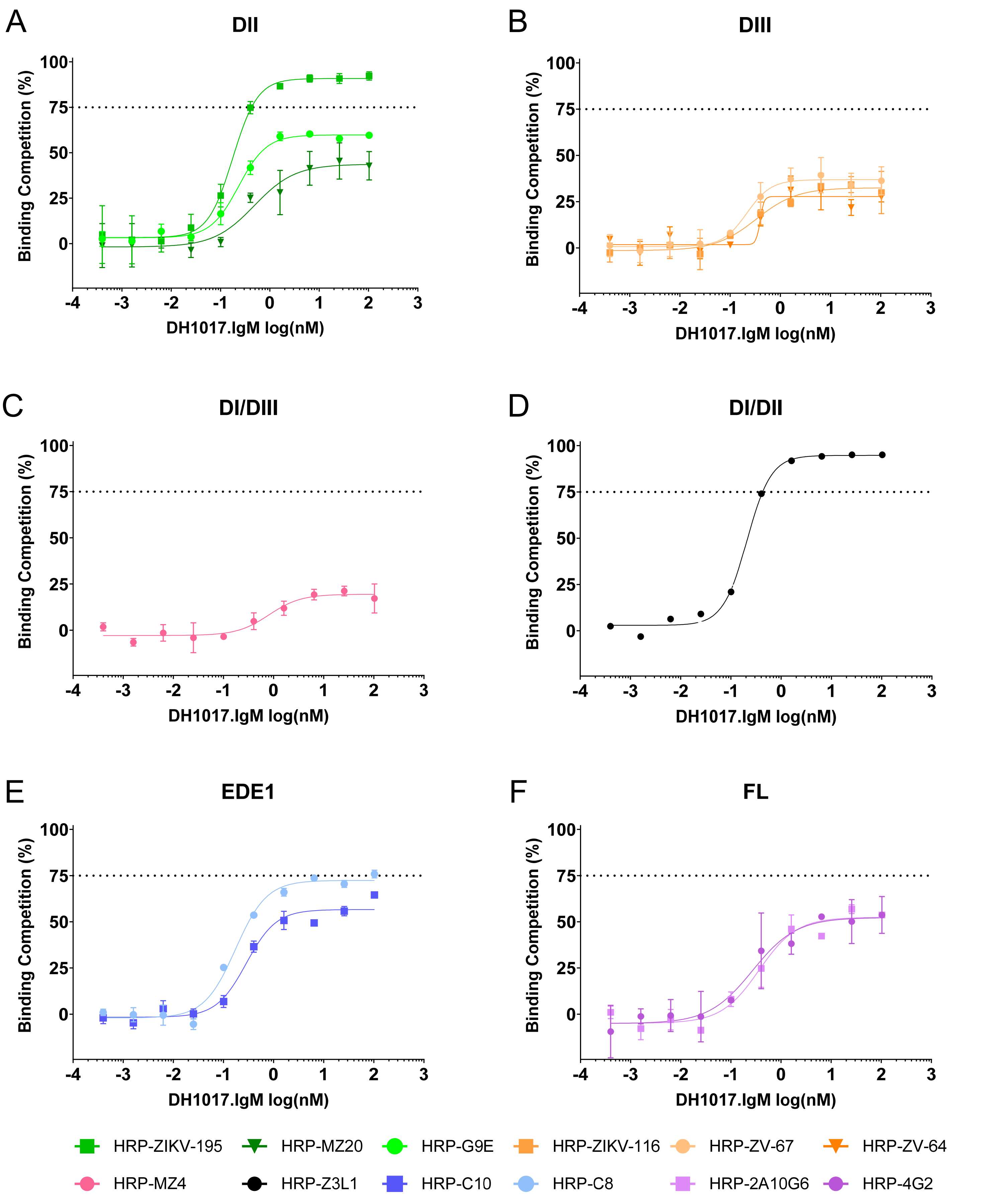


**Figure S3. DH1017.IgM dose-dependent competition of ZIKV-IgG mAbs binding to ZIKV-DAK whole virion.** Dose-response curves show the percentage of each of 12 ZIKV-IgG mAbs binding to ZIKV-DAK competed by DH1017.IgM grouped in panels based on IgG mAb binding regions as reported in Figure S2. Dotted lines: 75% binding reduction of each ZIKV-IgG mAb binding. Error bars: SD of two technical replicates. Data are representative of at least two independent experiments.

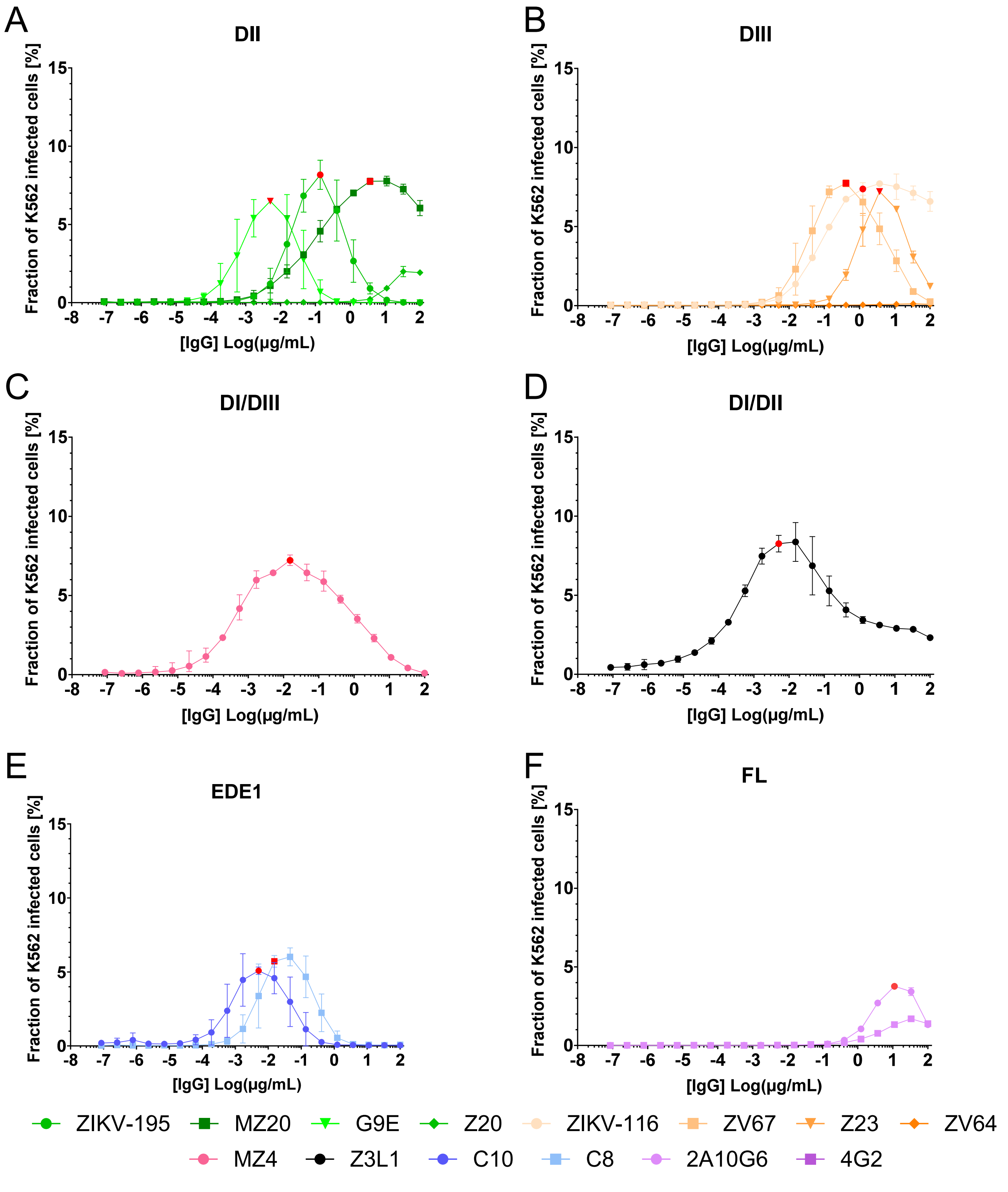


**Figure S4: Antibody-dependent enhancement of infectivity (ADE) profile of the ZIKV-IgG mAbs.** ADE-mediated infection of GFP-expressing ZIKV-DAK RVPs are reported for each of 14 ZIKV-IgG mAbs grouped in panels based on the respective binding regions as reported in Figure S2. The y-axis shows the percentage of infected FcγR-expressing K562 cells over the total number of acquired cells. Red dots: IgG mAb concentrations at peak ADE selected for subsequent analyses; when no clear ADE peak concentration could not be identified, the lower concentration with maximum ADE infectivity was selected. Error bars: SD of two independent replicates.

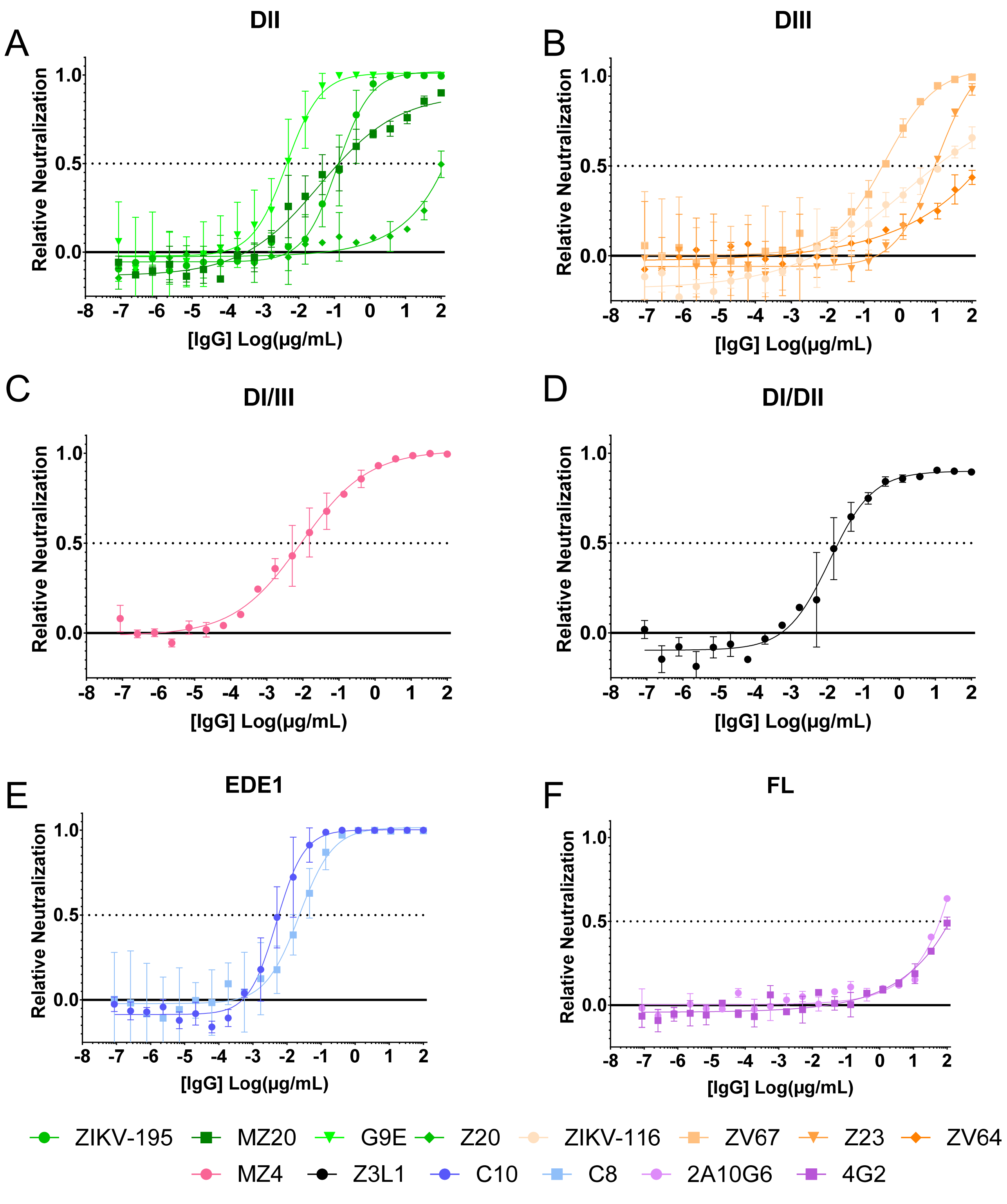


**Figure S5: Neutralization profile of the ZIKV-reactive IgG mAbs.** Neutralization of GFP-expressing ZIKV-DAK RVPs on Raji DC-SIGNR cells are reported for each of 14 ZIKV-IgG mAbs grouped in panels based on the respective binding regions as reported in Figure S2. The y-axis shows neutralizing activity relative to the medium-only condition. Dotted line: 50% neutralization ($RVPNC_{50}$), Error bars: SD of two independent experiments.

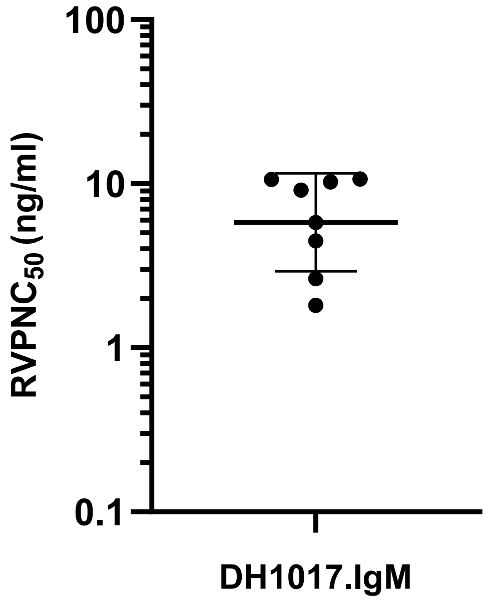


**Figure S6: DH1017.IgM neutralization potency against the ZIKV-DAK RVP strain.** $RVPNC_{50}$ of DH1017.IgM of GFP-expressing ZIKV-DAK RVP on Raji DC-SIGNR cells from 8 independent experiments. Line: Geometric mean; error bar: geometric SD.

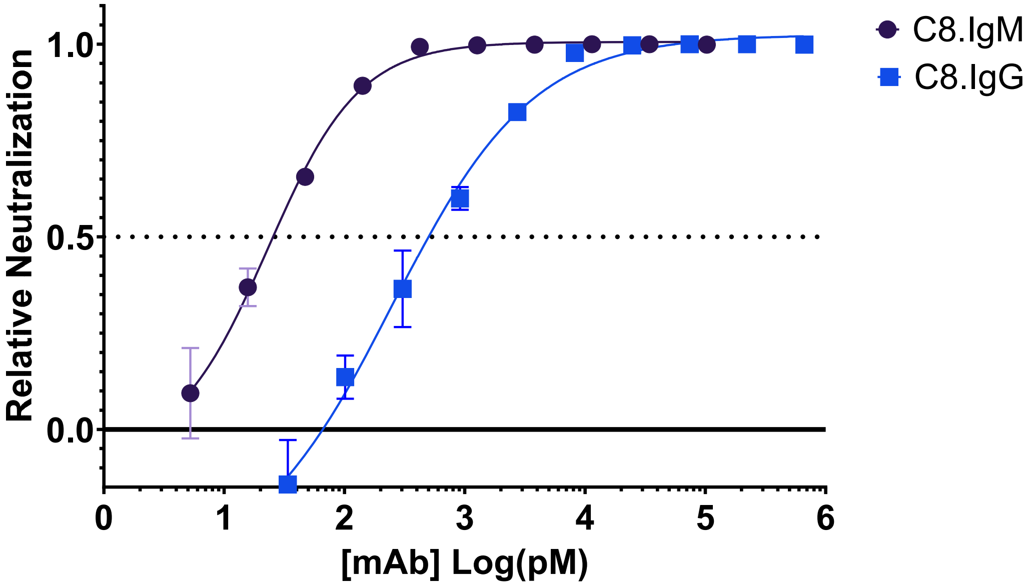


**Figure S7: Neutralizing activity of C8.IgM and C8.IgG.** C8.IgG (blue) and C8.IgM (black) dose-dependent neutralization of GFP-expressing ZIKV-DAK RVPs infectivity of Raji DC-SIGNR cells are reported relative to the medium-only condition. Concentrations are reported as pM with canonical molecular masses of 150 kDa and 970 kDa for the IgG and IgM isotypes, respectively. Dotted line: 50% neutralization ($RVPNC_{50}$), Error bars: SD of two independent experiments.

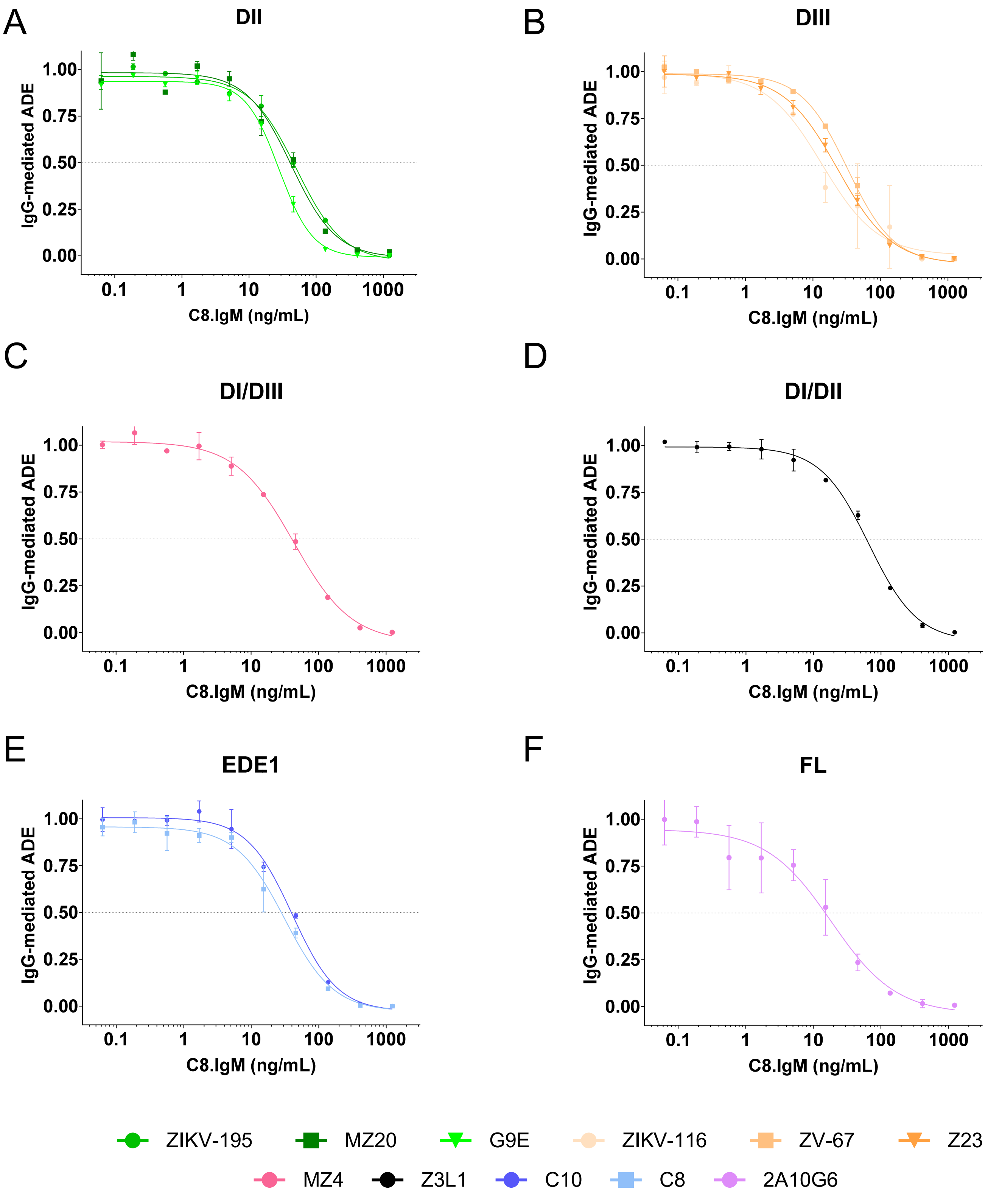


**Figure S8: C8.IgM dose-dependent potency in reducing ZIKV-IgG mediated ADE infectivity.** Each panel shows ADE-mediated infection of GFP-expressing ZIKV-DAK RVPs by a ZIKV-IgG mAb at their respective peak ADE concentration in presence of serially diluted C8.IgM. The y-axes show the C8.IgM dose-dependent reduction in ADE relative to the ZIKV-IgG alone condition. ZIKV-IgG mAbs are grouped in panels based on binding regions: **(A)** DII, **(B)** DIII, **(C)** DI/DIII linker, **(D)** DI/DII hinge, **(E)** EDE1, and **(F)** fusion loop (FL). Error bars: SD from two independent experiments. Dotted line: 50% reduction of ADE-infectivity ($cADEC_{50}$).
